# The ratio of RUNX1-ETO oncoprotein to normal RUNX1 expression determines the balance between endothelial reprogramming and hematopoietic cell growth

**DOI:** 10.64898/2025.12.19.695648

**Authors:** Floyd Hassenrueck, Katerina Terolli, Alexander Maytum, Peter Keane, Lea Flippe, Sophie Kellaway, Edouard G. Stanley, Elizabeth S. Ng, Andrew G. Elefanty, Constanze Bonifer

**Affiliations:** Murdoch Children’s Research Institute, The Royal Children’s Hospital, Parkville, Victoria 3052, Australia; School of Biosciences, College of Life and Environmental Sciences, University of Birmingham, UK, B15 2TT; Institute of Cancer and Genomic Sciences, School of Medicine and Dentistry, University of Birmingham, UK, UK, B15 2TT; Blood Cancer and Stem Cell Group, University of Nottingham, UK, Nottingham, NG7 2RD; The Novo Nordisk Foundation Center for Stem Cell Medicine (reNEW), Murdoch Children’s Research Institute, Parkville, Victoria 3052, Australia; Department of Paediatrics, Faculty of Medicine, Dentistry and Health Sciences, University of Melbourne, Parkville, Victoria 3052, Australia

## Abstract

In t(8;21) acute myeloid leukemia (AML) the RUNX1 DNA binding domain is fused to the RUNX1T1 protein producing the RUNX1-ETO onco-fusion protein. We previously showed that t(8;21) leukemic stem cells aberrantly activate endothelial signalling pathways to initiate blast cell proliferation. Here, we employed a human embryonic stem cell (ESC) line expressing an inducible RUNX1-ETO transgene to determine whether RUNX1-ETO directly induces endothelial signalling pathways or whether it is dependent on AML progression. Using single cell analyses we show that RUNX1-ETO induction reprograms ESC-derived myeloid progenitors towards endothelial cells. Integrated data analysis demonstrates (i) that endothelial reprogramming is RUNX1-ETO concentration-dependent and irreversible, (ii) that when RUNX1-ETO and RUNX1 expression levels are equal, a sub-population of blood progenitors escapes reprogramming and (iii) keeps proliferating when co-cultured with endothelial cells. Our experiments provide important insights into the earliest stages of epigenetic reprogramming by oncogenic transcription factors in AML.

## Introduction

Acute myeloid leukemia (AML) is caused by multiple mutations which affect transcriptional regulation by altering the function of transcription factors (TFs), chromatin modifiers or signalling molecules, impacting on hematopoietic differentiation. AML driven by the t(8;21) chromosomal translocation is one of the best characterised subtypes^1^. The translocation produces the RUNX1-ETO oncoprotein by fusing the DNA binding domain of RUNX1 to the RUNX1T1 (ETO) protein. Transcription of the fusion gene encoding RUNX1-ETO is regulated by the *RUNX1* promoters and is therefore active at the same developmental stages as the wild type *RUNX1* gene. Importantly, RUNX1-ETO interferes with the normal action of RUNX1 by binding to the same binding motifs in the genome ^2–4^. Similar to the phenotype observed in *Runx1* knock out mice, germline expression of RUNX1-ETO results in embryonic lethality with a complete lack of fetal liver hematopoiesis ^5–8^. Mice expressing a transgene encoding full length RUNX1-ETO in hematopoietic cells show a slowly evolving perturbed hematopoietic differentiation with expansion of the HSC compartment and the eventual development of an indolent MPD-like disease ^9,10^. This phenotype supports the conclusion that expression of RUNX1-ETO alone is not sufficient to cause AML, and that additional co-operating mutations are required for oncogenic transformation ^11^ ^12^. The majority of these are in genes encoding signalling molecules, such as constitutively activating mutations in the stem cell factor (SCF) receptor, KIT ^13^,.

An important feature of the pathology of t(8;21) AML is that the translocation can be found from an early stage of embryonic development onwards ^14^. Indeed, several studies have demonstrated the importance of the developmental stage at which the translocation occurs. Hematopoietic stem cells (HSCs) and their immediate precursors arise from the dorsal aorta of the vertebrate embryo via a process called the endothelial-hematopoietic transition (EHT) whereby cells bud off and move to the fetal liver and from there to the bone marrow ^15^. The EHT process involves (i) the upregulation of *RUNX1* in hemogenic endothelial cells, (ii) the activation of a blood cell specific gene regulatory network (GRN) by reshaping the chromatin and transcription factor binding landscape and (iii) the repression of the endothelial GRN by the concerted action of RUNX1 and interacting co-repressors ^16,17^. If RUNX1-ETO is expressed prior to the EHT, transition is blocked, and no blood cells are formed at all ^18,19^. If it is expressed after the EHT, it generates cell populations with enhanced self-renewal and reduced differentiation capacity^18,19^. However, since the translocation is a clonal event in vivo, it is still unclear how such cell populations develop.

An important consequence of leukemogenic mutations affecting gene regulation is the ’derailing’ of differentiation resulting from compensatory cellular changes in GRNs. This process can result in the ectopic activation of genes unrelated to myelopoiesis or even to blood cell development in general ^20–22^. Each AML mutational sub-type therefore displays a unique cellular identity and chromatin landscape which in t(8;21) AML is already manifest in leukemic stem cells (LSCs). Such cells maintain expression of the fusion transcripts and are quiescent, potentially serving as a reservoir for relapse of the disease. We recently showed that t(8;21) LSCs transiently reactivate aberrant signalling pathways to give rise to rapidly proliferating blast cells ^23^. One of these ectopically activated pathways is specific for endothelial cells and includes the gene encoding Vascular Endothelial Growth Factor alpha (VEGFA) and its receptor FLK1 which is encoded by the *KDR* gene ^23^. We also identified a molecular circuitry consisting of RUNX1-ETO, RUNX1, GATA2 and the MAP Kinase responsive transcription factor AP-1 as being responsible for LSC growth activation. However, how the expression of RUNX1-ETO activates this circuitry and how activation determines the phenotype of the cells directly after the translocation event is unknown.

To understand disease progression, it is therefore essential to study the immediate effects of the first mutational hit on the pathway to leukemogenesis. Moreover, mutations in genes encoding signalling proteins such as growth factor receptors that create constitutively active molecules usually occur after gene regulatory mutations, because their activation affects HSC self-renewal ^24^ ^25^. It is therefore necessary to separately dissect the contribution of each component to leukemogenesis. The differentiation of pluripotent stem cells (PSCs) into blood which recapitulate early hematopoietic specification including the EHT has been a valuable model enabling us to address such questions. We previously described a human PSC line which expressed an inducible version of RUNX1-ETO and showed that the induction of the oncoprotein led to extensive alterations of gene expression and chromatin structure ^18^. However, the precise phenotype of the induced cells at the single cell level was unknown.

To investigate the activation of aberrant signalling pathways resulting from RUNX1-ETO expression, we used an improved in vitro differentiation system capable of generating multi-lineage engrafting, definitive hematopoietic stem cells (iHSCs) followed by single cell analyses ^26^. RUNX1-ETO expression in myeloid precursors derived from iHSCs led to a (i) a dosage-dependent cell cycle block, and (ii) the rapid down-regulation of hematopoietic genes, both of which are associated with the loss of RUNX1 activity. Finally, RUNX1-ETO binding was correlated with an irreversible activation of an endothelial cellular phenotype responsive to VEGFA-mediated activation of MAP-Kinase signalling. However, when RUNX1-ETO expression equalled those of RUNX1, a sub-population of myeloid blood progenitors escaped endothelial reprogramming and grew within an endothelial environment. Our experiments validate the usefulness of our advanced differentiation system to gain insights into the primary events in leukemogenesis and provide a mechanistic explanation for the activation of endothelial genes in t(8,21) AML and the appearance of proliferating pre-leukemic cells.

## Results

### Induction of RUNX1-ETO in ESC derived myeloid cells leads to a dosage-dependent cell cycle block and activates an endothelial gene expression program

To dissect the molecular features of the pre-leukemic cells generated by RUNX1-ETO expression we examined the consequences of RUNX1-ETO induction in the context of our recently described PSC differentiation protocol that generates HSC-like cells, termed iHSCs ^26.18^ (Fig 1 A). After 13 – 15 days of differentiation, non-adherent CD34+ hematopoietic stem and progenitor cells (HSPCs) were expanded for up to eight days in a medium containing the hematopoietic cytokines SCF, TPO, IL-3, and EPO on an orbital shaker (Fig. 1A). RUNX1-ETO was induced by adding varying concentrations of Doxycycline (Dox) and the treated cells were subsequently characterised at different time points of induction using cell proliferation assays, surface marker and gene expression analyses. Relative levels of *RUNX1-ETO* and wild-type *RUNX1* mRNA expression were determined by qPCR.

**Figure 1.**
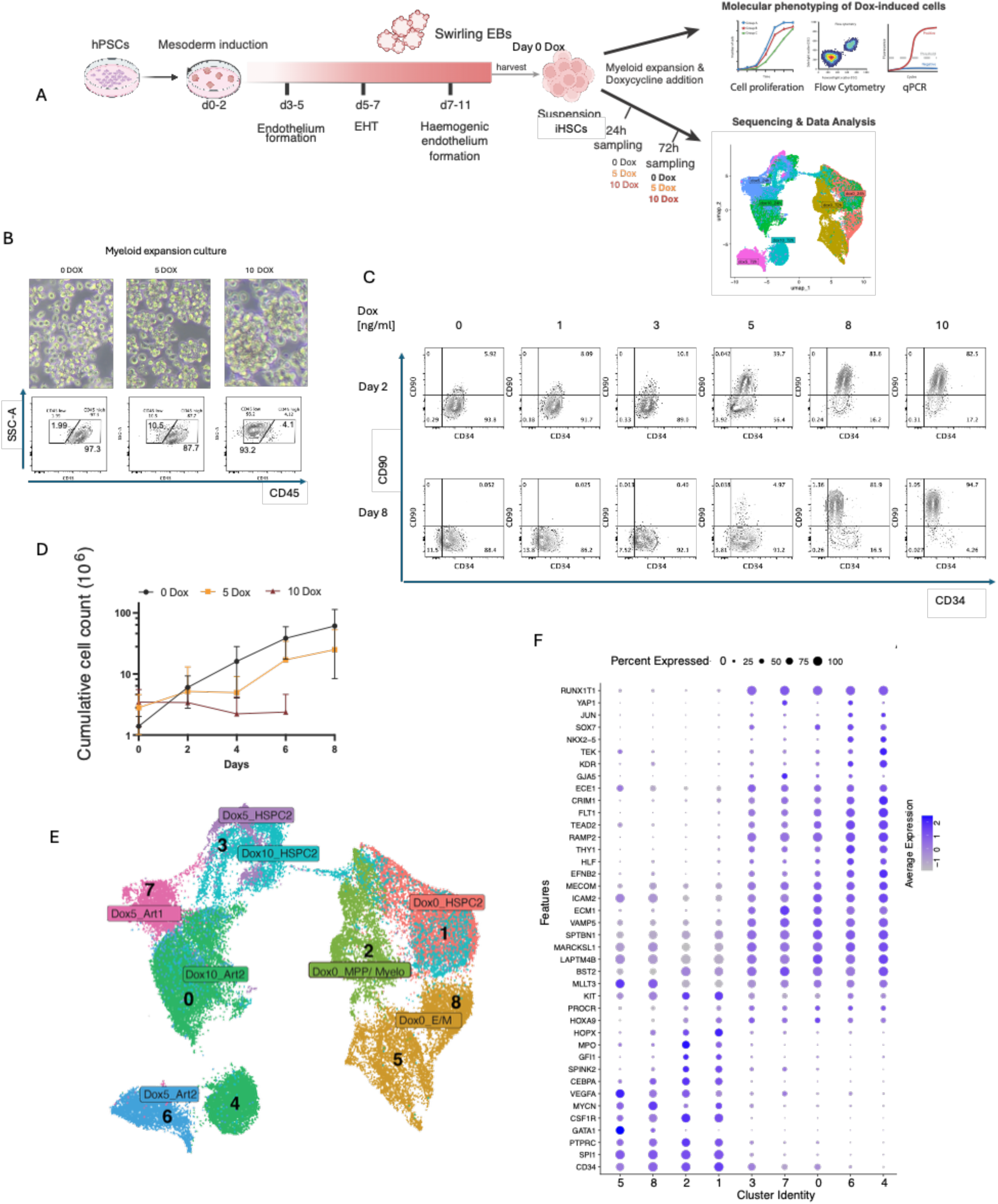
Expression of RUNX1-ETO leads to dosage-dependent arterial patterning and a cell cycle block. **A**: Schematic of the differentiation system showing the time course of hematopoietic differentiation from hPSCs to suspension iHSCs and further expansion after Doxycycline induction, followed by data analysis as depicted. Cells were expanded in rotating 6-well plates for up to 8 days and Doxycycline and growth medium were replenished every second day. Partially created using BioRender.com. **B.** Brightfield images of RUNX1-ETO HPSC derived hematopoietic cells under no doxycycline (0 Dox), 5 ng/ml Dox and 10 ng/ml Dox conditions showing formation of cellular aggregates loss of CD45 expression (bottom panel). **C.** Flow cytometric analysis of CD90 and CD34 surface marker expression at Day 2 and Day 8 of RUNX1-ETO induction using the indicated concentrations of Doxycycline (1,3,5,8,10 ng/ml) compared to uninduced cells (0 Dox). **D.** Box and whisker plot showing cumulative cell counts following Dox-induction (0,5,10 ng/mL) conditions over 8 days. Error bars, mean ±s.e.m., n=3 **E.** UMAP representation of 8 Seurat clusters highlighting cell populations with annotated hematopoietic and non-hematopoietic identity as identified in Supplemental Figure 1 and by comparison with embryo data ^26,27^. HSPC2: Hematopoietic stem and progenitor cells, embryonic cluster 2, Art1/2: Arterial cells, embryonic cluster 1 and 2, Myelo: Myeloid progenitor cells, E/M/M: Erythroid, megakaryocyte and mast cell progenitors. **F.** Dot plot of expression levels of differentially expressed genes in Seurat clusters representing arterial, hemogenic endothelium and hematopoietic genes.

The level of RUNX1-ETO expression was Dox concentration dependent between days 2 and 8 of induction but did not impact on the expression of *RUNX1* (Fig S1A, B). Within two days following the initiation of Dox treatment, cells dramatically altered their phenotype, changing from single non-adherent cells to cells forming large clusters (Fig 1B). Flow cytometry analysis showed this morphological change was accompanied by an increase in the side scatter of the treated cells (Fig. 1B, Fig. S1D), and coincided with loss of the hematopoietic surface marker, CD45 (Fig. 1C, Fig. S1C). At Dox concentrations of 5 ng/ml, at which point we have shown that the level of RUNX1 and RUNX1-ETO protein expression were equal ^18^, induction of RUNX-ETO resulted in the emergence of a CD34loCD90+ cell population (Fig. 1C). At higher Dox concentrations, cells displayed a profound, dosage-dependent growth impediment (Fig. 1D).

To gain insight into the nature of cells after RUNX1-ETO induction, we used single cell RNA-Seq analysis to examine the impact of induction time (24hr, 72hr) and Dox concentration (0, 5, 10 ng/ml) on altering gene expression in induced cells (Fig. 1A). We compared gene expression profiles to recently published single cell gene expression profiles of developing cells in the human embryo ^27^. This analysis had previously confirmed the HSC/HSPC phenotype of in vitro produced cells and identified cells of arterial origin giving rise to the hemogenic endothelium which then generated all hematopoietic lineages ^27^. The annotation of the different cell clusters arising in the myeloid expansion medium using the expression of cluster-defining genes (Fig 1F, Fig S1E, F, Fig S1I – M, Supplemental Dataset 1) revealed the presence of an HSPC cluster (Cluster 1) and a continuum of more differentiated myeloid (Cluster 2) and erythroid/megakaryocytic (Cluster 5,8) progenitor cells in uninduced cultures.

Induction of RUNX1-ETO led to dramatic changes in gene expression, with dox-induced cells forming distinct clusters from untreated cells including a Dox dependent cluster representing a subpopulation of cells with a cell cycle block in G1 (Fig. 1 E, Fig. S1 G, H). The annotation of the different clusters revealed that with the exception of cluster 3, which represented an intermediate cell type expressing a mixture of endothelial and hematopoietic genes that were mostly at the G2/M stage (Fig. S1 G), the majority of RUNX1-ETO expressing cells adopted an endothelial (arterial) phenotype, annotated Art1/Art2. Induced cells in clusters 0, 4, 6 and 7 expressed endothelial genes that were essentially absent from control cells as well as from clusters 1, 2, 5 and 8 that consisted of HSPCs (cluster 1) and differentiating early myeloid (cluster 2) and erythroid/megakaryocytic lineages (clusters 5, 8) (Fig. 1F). Pseudotime analysis performed by using tradeSeq^28^ (Fig 2 A) and Monocle^29^ (Fig. S2 A-C) showed (i) that all RUNX1-ETO expressing endothelial-like cells originated from HSPCs (cluster 1) and (ii) the reprogramming kinetics towards the arterial phenotype were Dox concentration and time dependent.

**Figure 2:**
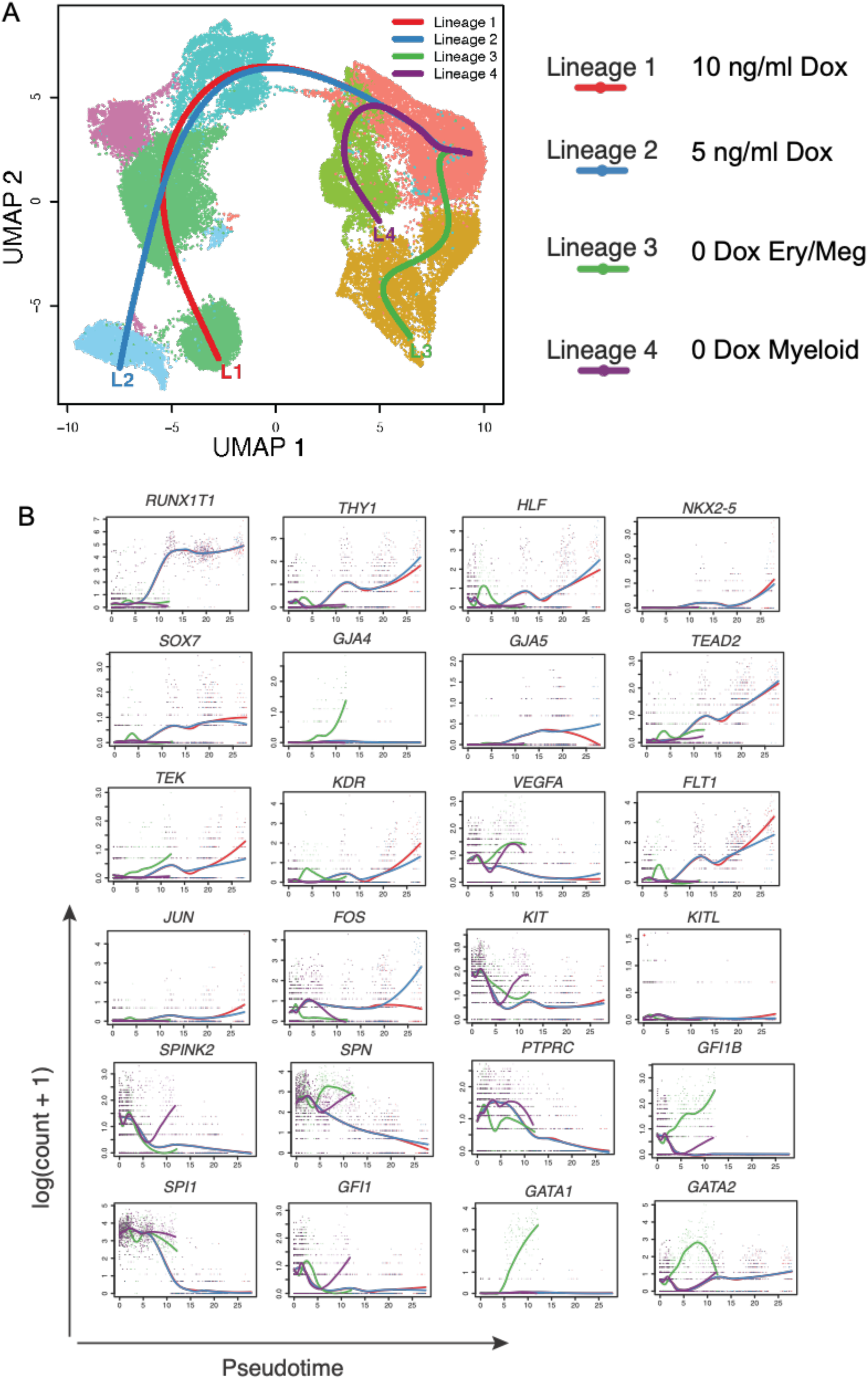
Expression of RUNX1-ETO alters the developmental control of gene expression and the differentiation trajectory. **A. :** Projection of the differentiation trajectory of cells in the four samples (0 Dox, 5 Dox and 10 Dox) on the 8 Seurat clusters highlighting cell populations with annotated hematopoietic and non-hematopoietic identity as in Fig 1E using trendSeq defining 4 lineages. **B. :** (0 dox, 5 dox and 10 dox) on the Seurat clusters using tradeSeq defining 4 lineages. Lineage 1, 10 dox; Lineage 2, 5 dox; Lineages 3 and 4, 0 dox.

Using tradeSeq to measure the number of cells expressing specific genes during a differentiation trajectory we next examined how the expression of marker genes for the different cell populations were dynamically changed during RUNX1-ETO induction (Fig 2B, Fig S2 D,E, Supplemental Datasets 2,3). When plotted against the different trajectories (1 – 4) detected in Fig 2 A, we noticed a synchronous induction of RUNX1-ETO (panel RUNX1T1) in all cells treated with either 5 or 10 ng/ml Dox verifying the cell-intrinsic nature of the biological effects we were observing. K-means clustering of co-ordinately regulated genes identified 4 with expression profiles that were specific for cells induced with 5 Dox (Fig S2 D, Supplementary dataset 2) and 5 clusters for unique to cells induced with 10 Dox (Fig S2E, Supplementary dataset 3), verifying the global nature of gene regulatory network shifts we observed.

The analysis of the dynamic regulation of specific genes showed a rapid down-regulation of the myeloid master regulator gene *SPI1* and the blood cell-specific *SPN* gene immediately after RUNX1-ETO induction (Fig. 2B). *GATA1* was exclusively upregulated in the erythroid/megakaryocyte (E/M) lineage in cultures not treated with Dox, accompanied by an initial transient upregulation in *GATA2* expression, which was then switched off as cells matured. *GFI1* and *GFI1B* were dynamically expressed in hematopoietic cells but were silenced following RUNX1-ETO induction in cells with an arterial phenotype. A mixture of HSC and endothelial genes expressed in the induced arterial cells such as *HLF, TEAD2, THY1, SOX7,* the angiopoietin receptor gene *TEK* and the VEGF-receptor genes *FLT1* and *KDR* paralleled the kinetics of upregulation of RUNX1-ETO suggesting a direct impact of the oncoprotein on their expression. An interesting Dox dose dependent expression pattern was seen in arterial cells representing the end of each trajectory, especially with the gene encoding the AP-1 component *FOS* ^30^. AP-1 mediates MAP Kinase signalling and mediates VEGF signalling via the VEGF-receptors ^23,31^. We observed a clear difference in *FOS* expression levels in response to the level of RUNX1-ETO induction, highlighting a difference in signalling responsiveness when RUNX1 and RUNX1-ETO are expressed at the same or at disparate levels.

### Induction of RUNX1-ETO perturbs the RUNX1 gene regulatory network

We next wished to explore how the binding of RUNX1 and RUNX1-ETO to chromatin affected the expression of genes associated with RUNX1 binding sites and to determine which cell populations were affected. RUNX1 can act as both a repressor of endothelial genes and as an activator of hematopoietic genes ^16,32,33^ whilst RUNX1-ETO has been shown to recruit co-repressors leading to repression of gene expression in transient transfection assays ^2,34,35^. We first determined the pattern and the dynamics of RUNX1/ RUNX1-ETO binding using our published ChIP data from hematopoietic progenitor cells representing a mixed population of immature and more differentiated HSPCs which had been either left uninduced or were induced with 5ng/ml Dox^18^. Examples of the dynamics of RUNX1 and RUNX1-ETO binding at specific genes are shown in Fig 3A and Fig S3 A,B). The *KIT* gene is a RUNX1 target whose expression is downregulated in endothelial cells after RUNX1-ETO induction. This reduction in expression coincides with a loss of RUNX1 binding at an intronic enhancer and the binding RUNX1-ETO to the RUNX1 promoter, leading to a reduction of RUNX1 binding (Fig 3A). Similarly, expression of *VAV1,* which shows RUNX1 binding at promoter and distal elements, is also diminished by RUNX1-ETO binding (Fig S3 A). Likewise, the *VEGFA* gene is bound by RUNX1 in uninduced cells, and its expression is reduced after binding of RUNX1-ETO at the same sites after induction (Fig S3B).

**Figure 3:**
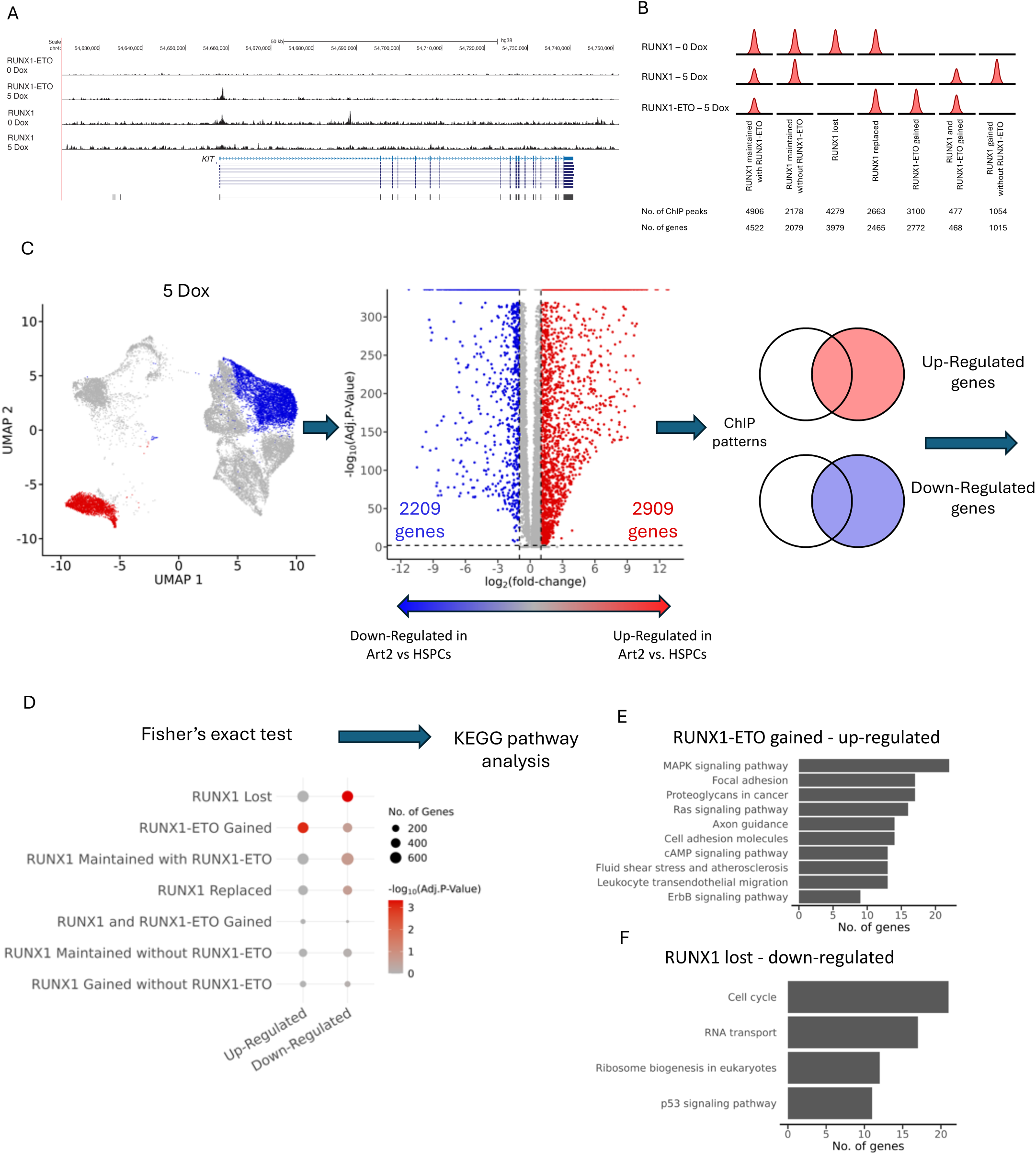
RUNX1-ETO interferes with the RUNX1-driven gene regulatory network. **A.** UCSC browser screenshots showing RUNX1 and RUNX1-ETO binding at the *KIT* locus in the presence or absence of 5 ng/ml Dox. **B.** Depiction of seven different classes on RUNX1 and RUNX1-ETO binding patterns obtained from previously conducted ChiP experiments using the same cell line ^18^ from uninduced or 5 Dox induced HSPCs. Peaks were filtered against ATAC-Seq peaks. Shared peaks are depicted with half the height. The table below summarizes peak numbers per group. Genes associated with binding sites as determined by comparing accessible chromatin peaks to those known to be associated with their rightful promoter as determined by comparison with promoter-capture HiC data from human CD34+ cells ^51^ are listed in a bottom line. **C.** Strategy of determining up- and down-regulated RUNX1 and RUNX1-ETO target genes in the 5 Dox-induced Art2 cluster (red) as compared to that of the uninduced HSPC2 cluster (blue). Left panel: UMAP of 0 and 5 dox cells highlighting the HSPC and Art2 cell populations at the start and end of the differentiation trajectory. Right: Volcano plot of genes which are differentially expressed between HSPC and Art2 cell populations in 0 and 5 dox cells. A positive fold-change indicates a gene was up-regulated in Art2 compared to HSPCs, and a negative value indicates a gene was down-regulated in Art2 compared to HSPCs. A gene was considered differentially expressed if it had a 2-fold change and an adjusted p-value less than 0.01. Right panel: Integration of differential gene expression values with the different patterns of RUNX1 and RUNX1-ETO binding as depicted in (B) **D.** Results from a one-sided (right tailed) Fisher’s exact test comparing the overlap of genes which were up and down-regulated in Art2 vs. HSPCs to ChIP binding groups depicted in Fig 2B. A significant p-value means that the number of genes which overlapped between groups was greater than could be expected by chance. Gene sets with a Benjamini-Hochberg corrected p-value < 0.05 was considered significant. **E, F.** KEGG pathway enrichment of gene sets that were found to be significant, in this case up-regulated genes gaining RUNX1-ETO binding (E) and down-regulated genes that have lost RUNX1 binding. Lists of enriched genes can be found in (Supplemental dataset 5)

Our global analysis shows that 2178 RUNX1 peaks were maintained but the majority of sites either lost RUNX1 binding after induction or binding was diminished as a result of RUNX1-ETO binding to the same sites (Fig 3B). We also counted 1054 RUNX1-only peaks, 4096 maintained but shared peaks and 3100 RUNX1-ETO-only peaks. Thereafter, we used published HiC data for human hematopoietic progenitor cells ^20,36^ to assign binding sites to genes (Fig 3B, bottom panel). We next sought to determine which genes were differentially expressed between cells from control versus Dox induced cultures. Using scRNA data, we compared gene expression within cells representing Cluster 1 (control) to gene expression within clusters 4 (5 ng/ml Dox) and 6 (10 ng/ml Dox) (Fig 3C, Fig S3C, middle panels) representing endothelial cells (Art2). This analysis was integrated with the ChIP data to gain an insight into the regulatory mechanisms underpinning the observed changes in gene expression. Specifically, we investigated whether changes in expression of certain groups of genes were associated with specific binding behaviour of RUNX1/RUNX1-ETO (Fig 3D, Supplemental datafile 5) using a Fisher’s exact statistical test (Fig 3D, Supplemental datafile 5). This analysis revealed that the up-regulation of genes in endothelial cells was correlated with the gain of RUNX1-ETO binding, whilst the down-regulation of genes in these cells correlated with the loss of RUNX1 binding at these genes as compared to HSPCs. Importantly, both enriched groups of genes affected different pathways (Fig 3E, F). The loss of RUNX1 affected the expression of cell cycle and ribosome biogenesis genes (Fig 3E, Supplemental datafile 5), which links this binding pattern to the loss of proliferation. RUNX1-ETO bound to up-regulated genes of the MAP-Kinase / RAS signalling and focal adhesion / angiogenesis pathway such as *JUN, FOS, FYN* as well as *VAV3* and *ITGA9* (Fig 3F, Supplemental datafile 5), linking this binding pattern to the activation of the endothelial phenotype.

### Induction of RUNX1-ETO blocks hematopoietic differentiation and generates endothelial cells in a dose-dependent fashion

We next studied the effects of RUNX1-ETO induction on the differentiation of hematopoietic cells, and whether the withdrawal of the Dox inducer restored the normal phenotype. To this end, we performed methylcellulose (MC) colony forming assays in the presence or absence of Dox as shown in Fig. 4A. We tested (i) whether the timing and duration of Dox treatment impacted the ability of cells form colonies, and (ii) whether these variables affected the nature of the colonies that were formed (Fig. 4 A-E). After 2 days of further culture under control conditions in the absence of Dox (0 Dox), HSPCs generated a similar frequency and spectrum of CFU-GEMM (colony forming unit - granulocyte, erythrocyte, macrophage, megakaryocyte), CFU-GM (granulocyte, macrophage) and CFU-G (granulocyte) colonies as day 15 cells (Fig. 4B,C). Treatment of cells with 5ng/ml Dox for 2 days prior to the initiation of colony forming assays did not significantly reduce the total colony number, when Dox was absent from MC cultures (5 Dox - Disc). However, maintaining Dox within MC cultures led to a significant reduction in the number of colonies (5 Dox - Cont) (Fig. 4B), reminiscent of the observed block in the cell cycle. A more pronounced effect was observed when cells were treated with 10 ng/ml Dox prior to MC assay initiation, with a significant reduction in the number of CFUs (10 Dox – Disc) in MC cultures lacking Dox and the complete absence of CFUs when Dox was maintained (10 Dox – Cont) (Fig. 4 B, C). Cells from cultures grown for 8 days the absence of Dox (0 Dox) generated no CFUs, suggesting these cells had differentiated beyond a clonogenic progenitor stage (Fig. 4 D,E). However, HSPCs treated with 5ng/ml Dox for 8 days prior to seeding into MC assays retained the capacity to form colonies, irrespective of whether the Dox treatment was continued (5 Dox – Cont) or withdrawn (5 Dox – Disc) (Fig 4. D, E). This retention of colony forming ability is consistent with RUNX1-ETO expression impairing myeloid differentiation. Under conditions in which RUNX1-ETO expression was continued during the MC assay no mature macrophage colonies (CFU-M) formed in contrast to those where Dox was discontinued. Similar to the results following a 2-day expansion, clonogenic cells were seen in 8-day expansion cultures when 10 ng/ml Dox was withdrawn prior to methylcellulose culture (Fig. 4 D, E).

**Figure 4:**
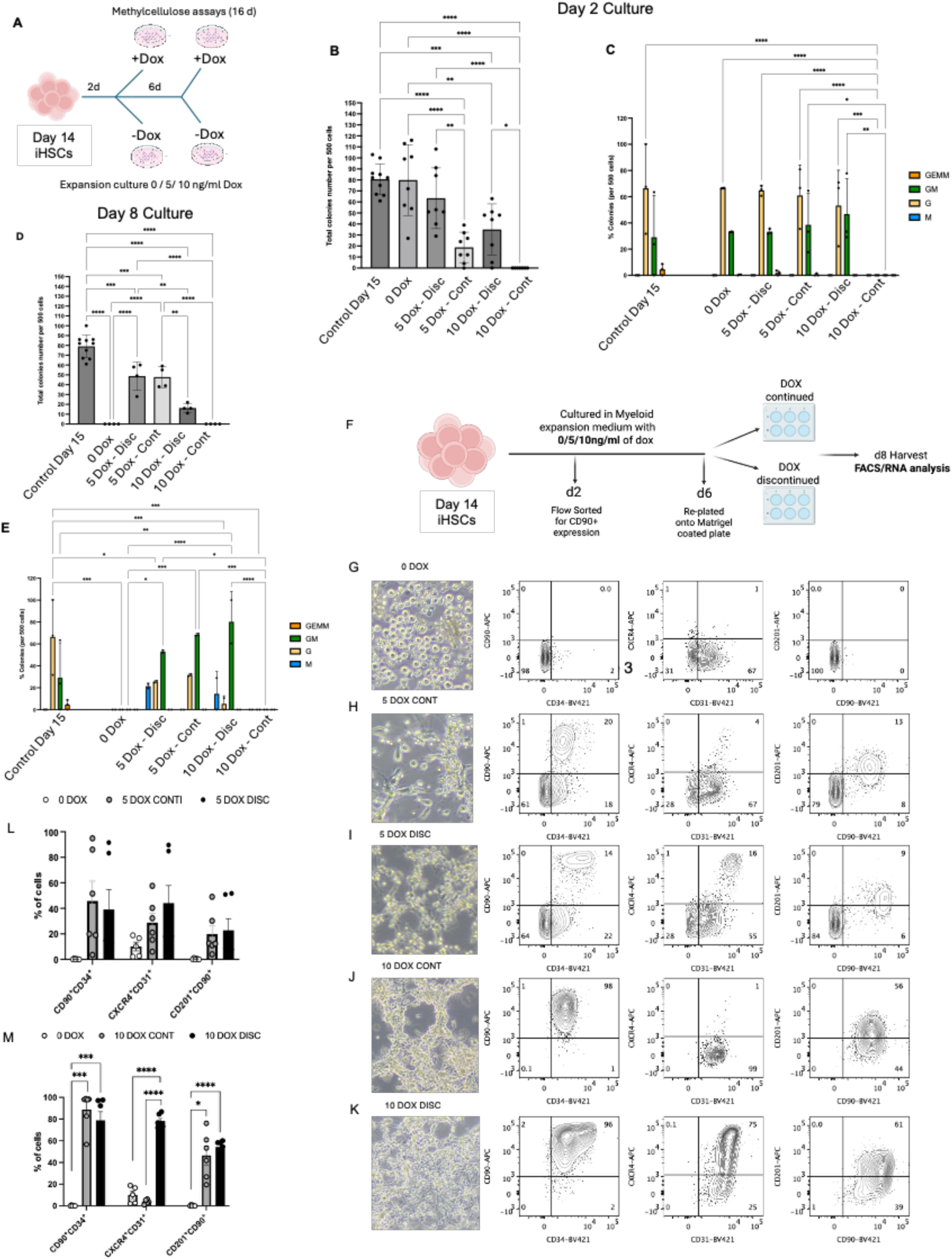
Induction of RUNX1-ETO blocks hematopoietic colony formation and generates endothelial cells in a dose-dependent fashion. **(A)** Scheme of colony assays. Day 15 iHSCs were sorted and cultured for 2 days in myeloid expansion medium with or without doxycycline (Dox), sorted on CD90^+^CD34^+^ expression and then plated in methylcellulose-based medium and cultured for 16 days in the presence or absence of Dox. Partially created using BioRender.com. **(B)** : Total number of colonies per 500 cells plated at day 2 of culture. 0 Dox, uninduced cultures; 5 Dox-Disc, 5 ng/ml dox induction and dox withdrawn from methylcellulose; 5 Dox-Cont, 5 ng/ml dox induction and dox continued in methylcellulose; 10 Dox-Disc, 10 ng/ml dox induction and dox withdrawn from methylcellulose; 10 Dox-Cont, 10 ng/ml dox induction and dox continued in methylcellulose. **(C)** Colony types quantified as CFU-GEMM (granulocyte, erythrocyte, monocyte, megakaryocyte), CFU-GM (granulocyte/macrophage), CFU-G (granulocyte), and CFU-M (macrophage), expressed as a percentage of total colonies found in (B). **(D, E)** Same as (B,C) but following 6 days of culture in expansion medium before methylcellulose plating. Bars represent mean ± SD from *n* = 3 independent experiments. Statistical significance was assessed using an Ordinary One-way ANOVA (B,D) or a 2way ANOVA (C,E) followed by Tukey’s multiple comparisons test. *p < 0.05; **p < 0.01; ***p < 0.001; ****p<0.0001. **(F)** Schematic of endothelial reprogramming and withdrawal experiments. Day 14 iHSCs were expanded in myeloid expansion medium with 0, 5 ng/ml or 10 ng/ml dox followed by flow sorting for CD90 and CD34 expression after 2 days. After 4 days further culture, cells were replated in endothelial assays in Matrigel with continuation or withdrawal of dox without addition of hematopoietic growth factors for 2 days. After a total of 8 days, cells were harvested for flow cytometry and transcriptomic analysis. Partially created using BioRender.com. **G – K.** Images (left panels) and flow cytometry analyses of cells after a total of 8 days culture (F) with the indicated surface markers. (**G**) uninduced cells (0 Dox ). **(H)** 5 Dox Cont, 5 ng/ml dox induction and dox continued in methylcellulose. **(I)** 5 Dox Disc, 5 ng/ml dox induction and dox discontinued in methylcellulose. **(J)** 10 Dox Cont, 10 ng/ml dox induction and dox continued in methylcellulose. **(K)** 10 Dox Disc, 10 ng/ml dox induction and dox discontinued in methylcellulose. Representative analyses of n=6 experiments are shown. **(L, M.)** Bar graphs of flow cytometry data for the indicated surface markers in cells treated as shown in (F). Error bars, mean ±s.e.m., n=6 experiments.

Whilst the induction of a gene expression program resembling that of arterial cells was an intriguing indication that iPSC-derived HSPCs adopted an endothelial identity, we wished to test whether they displayed other endothelial functions (Fig. 4F). We differentiated day 14 iHSCs in a myeloid expansion medium with or without 5 or 10 ng/ml Dox. On day 2 of expansion, induced cells were flow cytometrically sorted to enrich for CD90+CD34+ expression, and uninduced cells were sorted based on CD34 and CD45 expression, because they lacked surface CD90 (see Fig. 1C). Sorted cells were re-cultured in fresh medium with the same growth factors. At Day 6, we cultured uninduced cells and induced cells in a Matrigel-based endothelial network assay in the presence of VEGF (Fig. 4 F-K). Inspection of the cells after 2 days in Matrigel culture revealed a Dox-dependent alteration in cell shape and surface marker expression as compared to uninduced cells (Fig. 4 G, H, J, left panels). Cells started to adhere, interact and form a network of tubes. These changes were accompanied by the up-regulation of endothelial markers on the surface (CD90, CD201) (Fig. 4 G, H, J, right panels). Again, the proportion of cells undergoing these changes was dosage dependent. Cultures of cells induced with 5ng/ml dox contained a mixture of hematopoietic and endothelial cells (Fig. 4 H) whereas 10 ng/ml dox cells contained few residual blood cells (Fig. 4 J).

### RUNX1-ETO mediated endothelial reprogramming is irreversible

Our phenotypic analysis showed that the level of RUNX1-ETO expression affected the growth and the proportion of hematopoietic cells and endothelial cells after induction. RUNX1-ETO has been identified as a potential therapeutic target for siRNA therapy as leukemic cells have been shown to differentiate after its mRNA was depleted, demonstrating that even in the context of multiple mutations the presence of the oncoprotein was essential for leukemogenesis ^2,4,37^. However, RUNX1-ETO recruits epigenetic regulators such as histone deacetylases ^34,35^. Our phenotypic analysis therefore did not exclude the possibility of persisting, heritable gene expression changes. To this end, we withdrew Dox from the induced cells and measured surface marker expression as described in Fig. 4F, I, K). Following withdrawal, RUNX1-ETO expression was several hundred-fold reduced within 2 days (Fig. S4B). The quantitative analysis of FACS profiles (Fig. 4 L, M, Fig. S4C - F) demonstrated that the induction of RUNX1-ETO caused a profound shift of cellular identity from a hematopoietic towards an endothelial phenotype. However, the fraction of remaining CD45+SSC^low^ hematopoietic cells in the culture was dosage dependent and did not revert after Dox withdrawal, indicating irreversibility in the induced endothelial phenotype (Fig. S4G). Moreover, once RUNX1-ETO was withdrawn, cells moved closer to an endothelial phenotype by upregulating the endothelial markers CXCR4 and CD31 (Fig 4 I,K, right panels). The RUNX1-ETO inducible PSC line carries a reporter for the *SOX17* gene which is expressed in the arterial and hemogenic endothelia. Uninduced cells showed no or little expression of *SOX17*, which increased in a Dox dose-dependent fashion and persisted upon withdrawal, also suggesting that once cells had committed to the endothelial fate, this decision was irreversible (Fig. S4H).

Taken together, our data show that (i) RUNX1-ETO blocks hematopoietic differentiation which can resume after the cell cycle block was released by withdrawal of the Dox inducer, (ii) once endothelial cells are formed, the withdrawal of the oncoprotein does not reverse their phenotype, (iii) RUNX1-ETO induction affects differentiation of an early multipotent progenitor cell prior to the CFU-GM stage, which is in concordance with the finding that the viability of myeloid cells is not dependent on RUNX1 activity anymore ^38^, (iv) when expression of RUNX1 / RUNX1-ETO is equal a subpopulation of induced cells retains hematopoietic potential.

### RUNX1-ETO expressing myeloid progenitors in an endothelial environment escape the cell cycle block

We wanted to gain more insight into the gene expression of cells in the Matrigel cultures in which dox was continued or withdrawn, as shown in Fig. 4F. Initial qPCR assays examining the expression of endothelial and hematopoietic genes in these mixed populations total confirmed the dynamic response to Dox withdrawal seen in the flow cytometry data. Increased *DLL4, KDR* and *PECAM1* (CD31) expression indicated arterial patterning, but these analyses did not inform which cells responded (Fig. S4 I). Using scRNA-Seq we therefore explored whether endothelial and hematopoietic cells induced with 5 and 10ng/ml Dox retained an imprint of RUNX1-ETO oncogene exposure after withdrawal. We performed scRNA-Seq analyses of four samples: The starting cell population (day 14 HSPCs), day 22 Matrigel cultures containing uninduced control cells and day 22 Matrigel cultures with and without continuation of 5 ng/ml Dox for the last 2 days of culture (as shown in Fig. 4G/F). The composite analysis of all four samples showed that the samples from the uninduced day 14 and day 22 populations clustered separately, but that the 5 Dox continued and 5 Dox withdrawn day 22 cells predominantly clustered together as one large population together with two smaller but distinct populations (Fig. S4 J). Cell type classification yielded 13 clusters containing hematopoietic and endothelial cells (Fig. S4 K, Fig. S5 A, B, Supplemental Data File 6)

Individual analysis of clusters showed that uninduced cells had differentiated from HSPCs to granulocytes by day 22 (Fig. 5 A,B, Supplemental Data File 7, Fig. S4 J,K), whilst induced cells still contained a significant proportion of myeloid precursor cells (cluster 6 in Fig. S4 K), demonstrating that cell differentiation was blocked after RUNX1-ETO induction, as also seen in the colony assays (Fig. 4 A-E). The analysis of the cell cycle status of this cell population demonstrated that uninduced day 14 iHSCs cells included a higher proportion of proliferating cells than uninduced cells at day 22, consistent with their more mature differentiation state (Fig 5 C). In addition, this analysis together with the analysis of marker genes (Fig. 4D, Fig. S5 A, B, Supplemental Data File 8) confirmed that clusters 8 and 13 in Fig. S4 K were arterial endothelium (Art).

**Figure 5:**
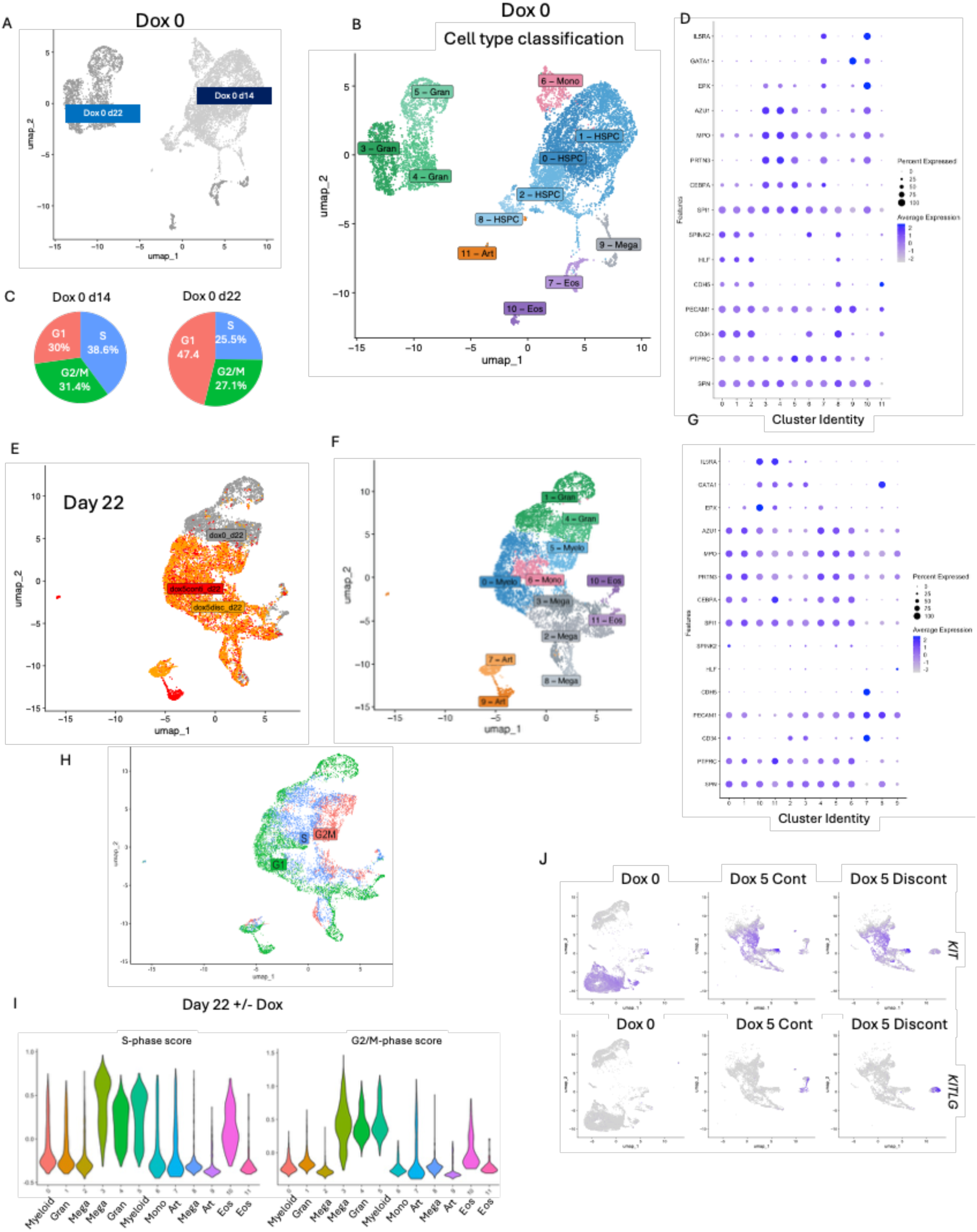
Endothelial reprogramming is irreversible, and endothelial culture supports the proliferation of RUNX1-ETO expressing hematopoietic cells. **A.** UMAP representation of scRNA-Seq data from uninduced cells at Day 14 and Day 22. **B.** Cell lineage annotation of uninduced cells at Day 14 and Day 22 to 11 clusters. HSPCs: Hematopoietic stem and progenitor cells; Art: Arterial endothelial cells; Gran: Granulocytic cells at different stages of maturation, Eos: Eosinophils, Mega: megakaryocytes, Mono: Monocytes. **C.** Cell cycle status of uninduced Day 14 and Day 22 populations. **D.** Dot plot of expression levels of differentially expressed genes in cells allocated to the Seurat clusters depicted in (B) representing stem cell and hematopoietic lineage genes. **E.** UMAP representation of scRNA-Seq data from Day 22 cells from Matrigel cultures that were uninduced or induced with 5ng/ml Dox, followed by Dox continuation (dox5conti) or discontinuation (dox5disc) for the final 2 days of Matrigel culture (see Fig. 4F). **F.** Cell lineage annotation of clusters of dox treated and untreated cells shown in (E) showing 9 clusters. Lineages as in (B). **G.** Dot plot of expression levels of differentially expressed genes in cells allocated to Seurat clusters depicted in (F) representing hematopoietic genes. **H.** UMAP representation of the cell cycle status of dox treated and untreated Day 22 cells from (E). **I.** S-phase and G2M- scores of dox treated and untreated clusters of Day 22 cells from (F). **J.** Feature plots from the same cell populations as in (F) separated by their induction status (Dox 0: No induction), continued induction with 5 ng/ml Dox (Dox 5 Cont) and after Dox withdrawal (Dox 5 Discont) showing the expression levels of *KIT* and *KITLG*

We directly compared uninduced day 22 Matrigel cultures to previously induced cells in which 5 ng/ml Dox was continued or withdrawn for the final 2 days culture (Fig. 4F, Fig 5 E – G), Fig. S5 C, Supplemental Data File 9). We found little difference in the gene expression profile between the different clusters with or without Dox withdrawal for the last 2 days which was also seen when feature plots of individual genes were examined (Fig. S5 C). The exception were the arterial clusters 7 and 9 IN FIG 5F, with cluster 9 representing cells in which 5ng/ml Dox was continued, and cluster 7 representing predominantly cells where Dox was withdrawn for the last two days (Fig. 5 E-G). Closer inspection of differentially expressed genes in these two cell populations (Fig. S5 D) showed multiple endothelial genes that were upregulated after Dox withdrawal, including CD34, CDH5 and CXCR4, which was also confirmed by our flow cytometry data (Fig. S4 C - F, Fig. 4 L, M). Withdrawal of dox upregulated a number of genes relevant for vascular biology such as the mechano-sensor *PIEZO2* ^39^ or *ROBO4* both of which are important for endothelial barrier function ^40^. These data suggested that cells progressed further towards endothelial differentiation without hematopoietic potential despite the withdrawal of Dox.

An interesting finding was that although our Matrigel cultures (Fig. 4 F-K) did not include any exogenously added hematopoietic specific cytokines, 5 Dox induced cultures contained a large proportion of hematopoietic cells even in the presence of RUNX1-ETO. This group of cells consisted of a heterogeneous population containing myeloid precursors, granulocytes, eosinophils and megakaryocytes (Fig. 5E-G). Furthermore, many of these cells were actively proliferating (Fig. 5 H) with a high S-phase score (Fig. 5 I). In addition, except for the arterial cells in clusters 7 and 9, we saw no difference in the gene expression profile in these cells (Fig. 5 F) in the presence and absence of Dox. This finding contrasted with the original 10 ng/ml Dox induction cultures which saw a large proportion of cells being reprogrammed and undergoing a cell cycle block, even in the presence of hematopoietic cytokines when the expression level of RUNX1-ETO exceeded that of RUNX1 (Fig. 1 D, Fig 4 F-K). However, when RUNX1-ETO and RUNX1 levels were equal a subset of hematopoietic cells continued to proliferate (Fig 1D). Endothelial cells are known to produce hematopoietic cytokines such as stem cell factor (SCF, encoded by the *KITLG* gene)^41^. Fig. 5 K shows that 5ng/ml Dox continued and withdrawn cells expressed the SCF receptor, KIT, but did not express *KITLG*. However, RUNX1-ETO induction leads to expression of this gene in the arterial cell clusters (7 and 9) even when Dox was withdrawn, (Supplemental dataset 9) suggesting that they could provide growth support for co-cultured hematopoietic cells. In addition, proliferating RUNX1-ETO induced cells express the VEGF receptor gene KDR (Fig. S1 L) which in VEGF containing cultures can drive the growth of leukemic myeloid progenitor and stem cells as seen in t(8;21) patients ^23^.

Taken together, our data show that RUNX1-ETO expressed at high levels acts as a roadblock to both blood cell division and differentiation, diverting cells to an apparently irreversible endothelial fate, but in the presence equal levels of RUNX1 and RUNX1-ETO, blood cells can harness available growth factor signalling to maintain growth.

## Discussion

Previous experiments using depletion approaches have identified a strong requirement in t(8;21) cells for the RUNX1-ETO oncoprotein for self-renewal, proliferation and the block of differentiation (Reviewed in reference ^1^). However, which of these features were a consequence of RUNX1-ETO expression and which ones were only acquired during disease progression has been unclear. Our study has shed light on the molecular details and the order of the earliest events during the transformation of myeloid progenitor cells by RUNX1-ETO, as summarized in Figure 6.

**Figure 6.**
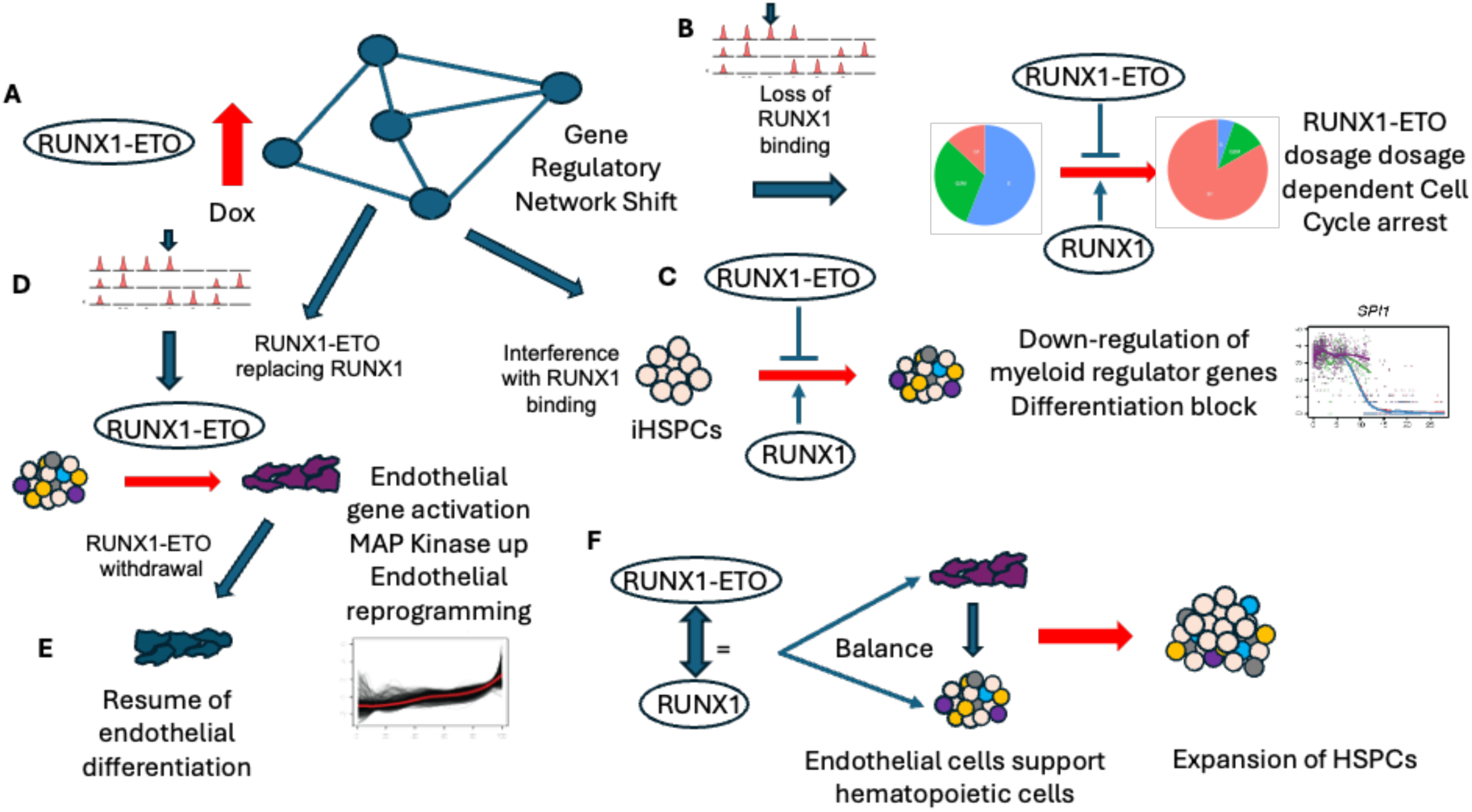
Model of early events in RUNX1-ETO initiated leukemia development. **(A)** RUNX1-ETO expression the gene regulatory network of myeloid progenitor cells and interferes with RUNX1 binding, **(B)** Deregulation of cell cycle genes is directly associated with RUNX1 loss and leads to a complete / partial cell cycle block, **(C)** Down-regulation of myeloid regulator genes and differentiation block seen with all concentrations of RUNX1-ETO **(D,E)** Rapid activation of endothelial gene expression program and MAP-Kinase pathway genes such as JUN directly associated with gain of RUNX1-ETO binding**, and (E)** Irreversible reprogramming of myeloid progenitor cells into endothelial cells after Dox withdrawal. **(F)** Partial maintenance of hematopoietic gene expression program when RUNX1 and RUNX1-ETO expression levels are equal, but impediment of myeloid differentiation. Co-culture of hematopoietic and endothelial cells bypasses the block in proliferation of blood progenitors in the absence of hematopoietic growth factors.

Once induced, RUNX1-ETO expression alters the gene regulatory network of myeloid progenitor cells and interferes with RUNX1 binding. The complete loss of RUNX1 leads to a deregulation of cell cycle genes and a cell cycle block, the degree depending on whether RUNX1-ETO is in excess or equal to that of RUNX1. A similar interplay with a requirement for balanced wild-type and mutated RUNX1 is evident in the context of several other Core Binding Factor (CBF) translocations ^42,43,44^. However, a wider requirement for RUNX1 expression in regulating growth and cell cycle progression is also seen in other types of AML such as the FLT3-ITD ^45^ ^46,47^. It is now clear that interference with RUNX1 activity leads to immediate consequences with regards to the growth of early progenitor cells.

However, the interference with growth is not the only effect of oncoprotein expression. After induction, the gain of RUNX1-ETO binding causes an activation of the endothelial gene expression program and of MAP-Kinase pathway genes such as *JUN* which irreversibly reprograms myeloid progenitor cells into endothelial cells in a dose dependent manner. Moreover, although the association with the binding pattern did not reach statistical significance, a large number of both up- and downregulated genes in arterial cells associate with a binding pattern that has RUNX1 and RUNX1-ETO binding to the same sites. The likely explanation for this finding is the ability of RUNX1 to act as both an activator and a repressor depending on the cellular context, with a major role in repressing endothelial gene expression ^16^. In addition, our finding provides a molecular explanation for the activation of endothelial genes seen in other RUNX1-related AMLs carrying the inv(16) or the t(3;21) translocations and CEBPA-double mutant AMLs which share the feature of a block of myeloid differentiation^44,47,48^.

Another surprising result was the irreversibility of endothelial cell reprogramming and the further progression of endothelial differentiation following RUNX1-ETO removal. In mouse embryos, the activation of RUNX1 in a RUNX1 null background was sufficient to rescue the hematopoietic potential of the hemogenic endothelium, but only at early developmental stages, providing supporting evidence for the idea that once endothelial differentiation has progressed, it is irreversible ^49^. RUNX1-ETO appears to halt this progress as well with cells sitting on the ’edge’ of two cell fates – endothelial and hematopoietic. An important result of our study with regards to the development of leukemic cells is therefore the appearance of sub-populations of cells maintaining the hematopoietic gene expression program when RUNX1 and RUNX1-ETO levels are equal, which are impaired, but not completely blocked in their differentiation. This result explains the reason for why such cells are still dependent on the presence of RUNX1. However, this feature was only evident during the co-culture of hematopoietic and endothelial cells which allowed hematopoietic cells to bypass the block in proliferation in the absence of exogenous hematopoietic growth factors even following the withdrawal of Dox induction of RUNX1-ETO. As Dox was only withdrawn for two days, we have yet to test whether such cells suffer from long-term effects of oncoprotein exposure with respect to their ability to differentiate as well. A recent study suggests that this could be the case, as it showed that depletion of RUNX1-ETO in patient cells the eliminated stemness features of leukemia initiating cells and differentiation from such cells was restricted to two lineages ^37^.

Several recent studies have shown that RUNX1 is intimately connected to growth factor signaling through its ability to cooperate with multiple signaling-responsive transcription factors, such as the mediators of MAP-Kinase signaling, the AP-1 factor family. In t(8;21) patient blast cells, AP-1 is required to maintain leukemic growth ^50^ and In t(8;21) leukemic stem cells, AP-1 mediates growth stimulation by VEGFA ^23^. In FLT3-ITD AML, RUNX1 chromatin binding is abolished in a subset of juxtaposed RUNX1-AP-1 binding sites when AP-1 binding is inhibited ^51^. Here we show that the binding of RUNX1-ETO leads to an immediate up-regulation of MAP-Kinase pathway genes suggesting that RUNX1-ETO induced cells are ready to rewire their growth factor responsiveness. However, t(8;21) patient cells down-regulate such genes including those driving VEGF-mediated growth once the oncoprotein is depleted ^23^, demonstrating that maintaining the right RUNX1/RUNX1-ETO balance is required throughout the entire process of disease progression.

t(8;21) patients accumulate secondary mutations that facilitate the growth of RUNX1-ETO expressing cells, such as mutations in KIT. Several studies, including those from our laboratory, have shown that it is possible to introduce a RUNX1-ETO expressing plasmid into human CD34+ HSPCs and obtain growing cells ^52,53^. The addition of a KIT-mutant expressing transgene further accelerates growth. However, as in our system, RUNX1-ETO expression levels need to be carefully adjusted. Moreover, many in vitro experiments grow cells in a sea of exogenously added cytokines including SCF, which is produced by the endothelial cells in our Matrigel cultures. It is therefore likely, that in contrast to a controlled induction of RUNX1-ETO cells, other culture systems may select for cells that can rapidly rewire their signaling needs to override the cell cycle block mediated by RUNX1-ETO.

In summary, our study has demonstrated that differentiated human pluripotent stem cells allow a faithful dissection of the initial stages of cellular reprogramming of hematopoietic cells by oncogenic transcription factors. Our study highlights the utility of pluripotent stem cells as a platform to screen for reagents aimed at therapeutic intervention.

## Limitations of this study

Like all in vitro studies, our experiments can only be an approximation of what occurs when a hematopoietic precursor cell undergoes a chromosomal translocation. The exact cell type that initially is mutated in vivo is not known, nor are the soluble and cellular components of the niche that comprise the milieu where this takes place. The Matrigel cultures demonstrate that signals from the arterial cells support hematopoietic cell growth but additional cells and signaling present in the hypoxic bone marrow niche may further support or antagonize growth. However, our observation provides a clear explanation how the RUNX1-ETO mediated cell cycle block can be bypassed and testing different hypotheses will be a major task in the future. The production of hematopoietic growth factors such as SCF is also seen in normal endothelial cells. Whether the arterial cells produced as a result of RUNX1-ETO expression are stable long-term in vivo and continue to impact on leukemic cells could not be addressed. We believe this to be unlikely given their reduced growth potential.

## Resource availability

### Lead contact

Further information and requests for resources/reagents/data should be directed to Constanze Bonifer or Andrew Elefanty

### Materials availability

This study did not generate new reagents. All cell lines can be shared upon request.

### Data and code availability

We re-analysed ChIP and ATAC datasets down-loaded from GEO from our previous publication ^18^ (GSE137673 and GSE137673, respectively). Single cell RNA-Seq data generated for this study have been deposited at GEO with the accession number **XXXXXXX**. Data are publicly available as of the date of publication. Any additional information required to reanalyse the data reported in this paper is available from the lead contact upon request.

Original new code has been deposited in https://github.com/Alexmaytum/RUNX1-ETO.Hassenrueck.Terolli.et.al/tree/main

### Author contributions

F.H., K.T and L.F. performed experiments and analyzed data, A.M., P.K. and C.B. analyzed data, S.K. provided data and edited the paper, E.G.S., E.S.N., A.G.E. and C.B. supervised the study, A.G.E. and C.B. wrote the paper. All authors approved the final version of the manuscript.

## Acknowledgements

We thank S. Rowley, Team Leader in the DNA Unit and Sequencing, Victorian Clinical Genetics Services, and her colleagues for sequencing of RNA samples, and M. Burton and E. Jones for flow cytometry assistance. This work was supported by the Novo Nordisk Foundation Center for Stem Cell Medicine, reNEW, supported by Novo Nordisk Foundation grant number NNF21CC0073729. This study was also funded by the National Health and Medical Research Council of Australia (NHMRC) through research fellowships GNT1117596 (A.G.E) and GNT1079004 (E.G.S.) and grants awarded to A.G.E. and E.G.S. (GNT1068866 and GNT1129861), to E.S.N. (GNT1164577 and GNT2012936), to A.G.E. (GNT2012535) and to E.G.S. (GNT1186019). Funding is acknowledged by the Stafford Fox Medical Research Foundation and the Victorian government’s Operational Infrastructure Support Program and Australian government’s NHMRC Independent Research Institute Infrastructure Support Scheme.

## Declaration of generative AI and AI-assisted technologies in the writing process

During the preparation of this work, no AI-assisted technologies were used.

## EXPERIMENTAL METHODS

### RUNX1-ETO induction experiments

The RUNX1-ETO expressing human embryonic stem cell line used in this study has been described in ^18^. Hematopoietic stem and progenitor cells (termed iHSCs) were generated as described ^26^ and harvested from days 13 - 15 and cryopreserved in CJ2/10% DMSO cryopreservation medium ^54^. iHSCs were thawed and cultured in SPELS medium base ^26^ supplemented with 100ng/ml recombinant human (rh) stem cell fator (SCF), 100ng/mL rh thrombopoietin (TPO), 25 ng/mL rh interleukin(IL)-3 and 2U/mL rh erythropoietin (EPO). Cells were cultured on an orbital rotating platform in 6-well plates for up to 8 days and RUNX1-ETO was induced using between 0-10ng/ml of doxycycline (Dox). Culture medium, including dox, was replenished every 2 days, and cell samples were taken for growth analysis, flow cytometry analysis and mRNA expression analysis. For growth analysis, cells were harvested every two days and counted using an automated cell counter (Nexcelcom Biosciences) using Propidium Iodide/Acridine Orange staining for live/dead cell discrimination. Cells were analysed by flow cytometry (BD Fortessa).

### RUNX1-ETO withdrawal experiments and Matrigel assay

Day 14 iHSCs were cultured for 6 days in myeloid expansion medium and treated with either 0, 5 or 10ng/ml of dox, replenished every 2 days. On day 2, dox treated cells were flow cytometrically sorted for CD90+CD34+ expression. Since only induced cells expressed CD90, uninduced cells were sorted based on CD34+ expression.

After 6 days of myeloid expansion, cells were replated onto Matrigel coated 6 well plates where dox treatment was either continued or withdrawn for a further 2 days. Cells were maintained in SPELS base medium supplemented with 2ng/ml rh bone morphogenetic protein (BMP), 5ng/ml rh insulin-like growth factor (IGF)1 and rh IGF2, 10ng/ml rh fibroblast growth factor (FGF) 2, 5ng/ml rh epidermal growth factor (EGF) and 25ng/ml rh vascular endothelial growth factor (VEGF). Cells were harvested at day 8 by gently aspirating floating cells followed by rinsing the well with SPELS. 1ml of 2ng/ml Dispase was added to each well and incubated for approximately 15 minutes at 37°C on a rotating platform at 60rpm. The wells were then flushed using SPELS to lift adherent cells. Harvested cells were analyzed using flow cytometry and qPCR.

### Cell sorting and flow cytometry

Cell surface marker expression of hematopoietic cells was analysed by flow cytometry using a BD LSRFortessa flow cytometer. Briefly, cell pellets were stained with antibodies for 15 minutes in the dark and washed with FACS wash buffer before resuspending in wash buffer containing propidium iodide to identify viable cells.

Hematopoietic cells were also single cell sorted for CD90 and CD34 cell surface expression using the BD FACSAria. A complete list of antibodies used for flow cytometry can be found in the STAR methods table.

### Methylcellulose Colony-forming assays

Hematopoietic colony forming cells in RUNX1-ETO induced and uninduced cultures were identified by culturing 500 cells in serum free methylcellulose, supplemented with 100ng/ml rh TPO, 100ng/ml rh SCF, 25ng/ml rh IL3, 25ng/ml rh IL6, 25ng/ml rh FLT3-ligand (FLT3L), 25ng/ml rh granulocyte colony stimulating factor (G-CSF) and 5U/ml rh EPO. Methylcellulose medium was prepared by mixing 40 mL of serum-free MethoCult™ H4100 with an equal volume of IMDM 2X supplement. The IMDM 2X supplement contained IMDM with 2x ITS-E, 0.2% rh albumin (Sartorius), 0.03% monothioglycerol (MTG), 0.2x SyntheChol, 125 ng/mL linolenic and linoleic acids, 1 µg/mL soybean oil, 2 mM GlutaMAX-I, 50 µg/mL L-ascorbic acid 2-phosphate (AA2P), and 50 µg/mL ascorbic acid. To this mixture, 20 mL of SPELS medium was added to achieve a final volume of 100 mL, along with 5 µM hydrocortisone and 9 ng/mL human low-density lipoprotein (hLDL). Cells were cultured in methylcellulose in the presence or absence of 5 or 10 ng/ml Dox. Dox was added to the methylcellulose every two days. Cultures were set up in duplicate in low attachment 24-well plates. Plates were scored for total hematopoietic colonies after 6 days and for colony subtypes after 16 days.

### RNA extraction and real time quantitative PCR analysis

Total RNA was extracted from cell samples using the Isolate II RNA Micro Kit (Bioline) following the manufacturer’s instructions. Complementary DNA (cDNA) was synthesized from purified RNA using random hexamer priming and the the Bioline Tetro cDNA Synthesis Kit (Bioline). Quantitative PCR (qPCR) was performed using cDNA as a template in combination with TaqMan Fast Advanced Master Mix (Thermo Fisher) and the TaqMan Probes described in the Key Resources table in a MicroAmp™ 96-well optical reaction plate. *GAPDH* (Hs99999905_m1) expression was used as a reference control. Reactions were run on a QuantStudio™ 5 Flex Real-Time PCR System (Applied Biosystems).

## SEQUENCING AND BIOINFORMATICS METHODS

### Single-Cell RNA Sequencing

#### Sample generation

Single-cell suspensions were generated from the different cultures and cells were fixed using the 10x Genomics Fixation Kit for Chromium Fixed RNA Profiling (CG000478 Rev C) according to the manufacturer’s instructions. Cell viability was assessed using a Nexcelcom Biosciences cell counter with acridine orange/propidium iodide (AOPI) staining. All samples were fixed overnight for 24 hours at 4°C and subsequently stored at –80°C until library preparation.

Samples were barcoded using the multiplexed human transcriptome kit for fixed RNA (10x Genomics PN-1000476), allowing sequencing of four samples in a single GEM reaction. Libraries were prepared using the 10x Genomics Chromium Controller, targeting 6,000 cells per sample. Barcoded single-cell cDNA libraries were generated and amplified according to the manufacturer’s workflow. Libraries were sequenced on an Illumina NovaSeq 6000 platform with paired-end reads to a depth of approximately 50,000 reads per cell.

#### Data Processing and Clustering

Raw base call files were converted to FASTQ format and aligned to the GRCh38-1.2.0 reference genome using Cell Ranger (v7.2.0) with default parameters. Cell barcodes and unique molecular identifiers (UMIs) were filtered and counted to generate gene expression matrices.

Downstream analyses were performed in R (v4.4.1) using Seurat (v5.1). Cells with fewer than 2,500 detected genes or more than 5% mitochondrial transcripts or more than 1×10^5^ transcripts were excluded. Gene expression matrices were log-normalized. Dimensionality reduction was performed using principal component analysis (PCA) followed by UMAP for visualization. Clustering was conducted using a shared nearest neighbour graph-based approach with resolution set to 0.3. Following initial clustering only those cluster containing more than 100 cells were included in downstream analyses.

Differential gene expression was assessed using the FindMarkers function of Seurat. Cell types were annotated based on canonical marker gene expression and confirmed by reference-based annotation using scType and comparison to known markers of HSPC subtypes as shown in ^26,27^.

#### Single-cell trajectory analysis

The processed single-cell expression data object from Seurat was imported into Monocle3 v1.4.26 ^55^ using the as.cell_data_set function from SeuratWrappers (https://github.com/satijalab/seurat-wrappers). Trajectories were then inferred using the learn_graph command in Monocle with the parameters close_loop = FALSE and use_partition = FALSE. Pseudotime was then calculated using the order_cells function using earliest principal node in the HSC population as the root node. These were then plotted using the UMAP coordinates calculated by Seurat.

#### Classification of RUNX1 and RUNX1-ETO binding patterns from ChIP-Seq data

ChIP-Seq data were down-loaded from GSE137673 and re-processed. Peak sets representing distinct RUNX1 and RUNX1-ETO binding patterns were defined using a custom python script (see code availability section). To do this, BED files from ChIP-Seq experiments were first filtered to keep only peaks found in open chromatin by comparing them to the union of all ATAC-Seq peaks using the intersect function in the pybedtools package v0.11.0 ^56^ in python v3.12.3. Only peaks that were found within an ATAC peak were kept for further analysis.

To ensure consistency in peak annotations between different ChIP-Seq experiments, peaks were mapped to a common set of coordinates by combining peaks from all experiments into a single pybedtools data object and merging them using the merge function. ChIP binding patterns were then classified by comparing the peaks from the RUNX1 0 dox, RUNX1 5 dox and RUNX1-ETO 5 dox samples to each other using the intersect function in pybedtools. The set of queries used to do this can be found in the *define_peak_sets.py* script (see code availability section). The resultant peak sets were then exported as BED files.

ChIP peaks were annotated to their target genes using *annotateBED_with_CHiC.py* script from Coleman *et al.* (2023) ^45^. This was done by comparing ChIP peaks in distal cis-regulatory elements to sites annotated to the promoter that they interact with using promoter-capture HiC datasets obtained from healthy PBSCs and t(8;21) AML patients. In cases where a peak could not be annotated with the HiC data, or was found at a promoter site, peaks were instead annotated to their closest gene using the annotatePeaks.pl function in HOMER v4.11 ^57^. The *annotateBED_with_CHiC.py* script and promoter-capture HiC annotation files used for this analysis can be found in the GitHub repository for Coleman *et al.* (2023) ^45^ at https://github.com/petebio/Gene_regulatory_network_analysis.

#### Comparison of ChIP-Seq binding patterns to changes in gene expression along the HSC to Art2 differentiation trajectory

Genes that change expression during Art2 differentiation after RUNX1-ETO induction were identified by selecting the single-cell clusters corresponding to earliest HSCs in the 0 dox sample and comparing these to the Art2 cells at the terminal end of the differentiation trajectory in either the 5 or 10 dox samples. This was done using the FindMarkers function in Seurat with the option min.pct = 0.1 to keep only genes that were detected in at least 10% of cells. A gene was considered to have significantly changed expression if it had a fold-change greater than 2 and a Bonferroni adjusted p-value less than 0.01.

To test if the set of up or down-regulated genes could be associated with specific changes in the pattern of RUNX1 and/or RUNX1-ETO binding, we compared them to the ChIP target gene sets associated with specific binding patterns. This was done using a one-sided (right-tailed) Fisher’s exact test in R v4.5.0 with the resulting p-values adjusted for multiple testing using the Benjamini-Hochberg method. Any comparison that had an adjusted p-value less than 0.05 was considered statistically significant.

KEGG pathway analysis was then conducted on the up or down-regulated ChIP target genes from significant comparisons using the enrichR package v3.4 ^58^ in R and using all genes that were expressed in at least 10% of cells in the single-cell clusters as the background gene set. Pathways with a Benjamini-Hochberg adjusted p-value less than 0.05 was considered statistically significant.

#### Analysis of gene expression along trajectories

Slingshot (version 2.16.0) ^59^ was used to infer trajectories on UMAP embeddings. Cluster 1 was set as the starting population and the clusters 5, 3, 4 and 6 were specified as end clusters. Slingshot was run with the commands extend= “n”, allow.breaks=TRUE, reweight=True, thresh=0.01 and maxit=30.

tradeSeq (version 1.22.0) ^28^ was then used to fit a negative-binomial generalised additive model for each gene along pseudotimes. 1500 cells sampled randomly were selected to be used in the model split equally across the lineages. The resulting set of cells were used to construct pseudotime matrices, lineage weight matrices and count matrices for tradeSeq analysis. Models were fit using nknots=9. associationTest was used to identify genes with differential expression along pseudotime.

For lineage specific pseudotime clustering, cells belonging to slingshot Lineage 1 (10 dox) and Lineage 2 (5 dox) were selected. Genes with less than 20 counts across the lineage were removed. Differentially expressed genes along the lineages were identified using tradeSeq and then filtered to include genes with padj<0.05 and which were in the top 40% ranked by absolute log fold-change. Smoothed expression trajectories were estimated across pseudotime for each gene using predictSmooth function. Gene by pseudotime matrices were then scaled by z-score scaled per gene.

To determine the number of clusters to use for K-means clustering ^60^, K-means was run for k=1-15 and the total within-clusters sum of squares was plotted to form an elbow plot. Final gene clusters were generated using R’s kmeans() function with centers=4 and nstart=25.

#### ATAC-seq Data Processing

Single-end ATAC-seq reads from sorted RUNX1c-positive and RUNX1c-negative populations were down-loaded from GSE137673 and were reprocessed by trimming using Trimmomatic (v0.39) ^61^ to remove low-quality bases and Nextera adaptors. Reads were aligned to the human genome (hg38) using Bowtie2 (v2.5.4) ^62^ with the --very-sensitive-local parameter. PCR duplicates were removed with Picard MarkDuplicates (v2.26.10). Peaks were called using MACS2 (v2.2.9.1) ^63^ with the parameters --nomodel -B --trackline and filtered against the hg38 ENCODE blacklist ^64^ using bedtools (v2.31.1) ^65^. Peaks were extended ±200 bp around the summit and annotated with HOMER annotatePeaks.pl ^57^.

#### ChIP-seq Data Processing

ChIP-seq reads were trimmed, aligned, and filtered as described above. Peaks were called with MACS2 using -B --tracklineand extended ±100 bp around the summit. ChIP-seq peaks were filtered against the merged ATAC-seq peaks from positive and negative RUNX1c populations to remove sites never accessible in ATAC-seq. Filtered peaks were annotated using the published annotate_bed_with_HiC pipeline ^45^, which integrates CHiC interaction data with HOMER’s annotatePeaks.pl to assign peaks to target genes.

#### Regulatory Element Classification

Annotated peaks were intersected across Dox0 and Dox5 RUNX1 samples and RUNX-ETO Dox5 samples using Intervene Venn ^66^ to define seven groups of regulatory elements as marked in Figure 2A. These groups formed the basis for integration with scRNA-seq trajectories.

#### Integration of regulatory elements with scRNA-seq trajectory

To assess the functional consequences of RUNX1 and RUNX-ETO binding on gene expression, the regulatory element groups were integrated with scRNA-seq pseudotime trajectories. Cells at the extremes of the pseudotime trajectory were selected to ensure accurate fold-change estimates and to minimize confounding from intermediate heterogeneous populations.

Differential gene expression between trajectory extremes was calculated using Seurat’s **FindMarkers** function, and genes were assigned to the seven groups based on prior ChIP-seq annotation. Gene set enrichment (GSEA, ^67^) and Leading-edge analyses were performed for each group to identify pathways and genes associated with distinct transcription factor binding patterns.

