## Supplementary Figures and Legends for "The ratio of RUNX1-ETO oncoprotein to normal RUNX1 expression determines the balance between endothelial reprogramming and hematopoietic cell growth": Hassenrueck, Terrolli et al., Bioarchive.pdf

**Resource availability**

**Materials availability:** This study did not generate new reagents. All cell lines can be shared upon request.

**Data and code availability:**

We re-analysed ChIP and ATAC datasets down-loaded from GEO from our previous publication [18] (GSE137673 and GSE137673, respectively). Single cell RNA-Seq data generated for this study have been deposited at GEO with the accession number **XXXXXXX**. Data are publicly available as of the date of publication. Any additional information required to reanalyse the data reported in this paper is available from the lead contact upon request.

Original new code has been deposited in https://github.com/Alexmaytum/RUNX1-ETO.Hassenrueck.Terolli.et.al/tree/main

**Author contributions**

F.H., K.T and L.F. performed experiments and analyzed data, A.M., P.K. and C.B. analyzed data, S.K. provided data and edited the paper, E.G.S., E.S.N., A.G.E. and C.B. supervised the study, A.G.E. and C.B. wrote the paper. All authors approved the final version of the manuscript.

**Acknowledgements**

We thank S. Rowley, Team Leader in the DNA Unit and Sequencing, Victorian Clinical Genetics Services, and her colleagues for sequencing of RNA samples, and

M. Burton and E. Jones for flow cytometry assistance. This work was supported by the Novo Nordisk Foundation Center for Stem Cell Medicine, reNEW, supported by Novo Nordisk Foundation grant number NNF21CC0073729. This study was also funded by the National Health and Medical Research Council of Australia (NHMRC) through research fellowships GNT1117596 (A.G.E) and GNT1079004 (E.G.S.) and grants awarded to A.G.E. and E.G.S. (GNT1068866 and GNT1129861), to E.S.N. (GNT1164577 and GNT2012936), to A.G.E. (GNT2012535) and to E.G.S. (GNT1186019). Funding is acknowledged by the Stafford Fox Medical Research Foundation and the Victorian government's Operational Infrastructure Support Program and Australian government's NHMRC Independent Research Institute Infrastructure Support Scheme.

### Declaration of interests

Andrew Elefanty, Elizabeth Ng and Edouard Stanley are consultants for Retro Biosciences Inc. The company had no involvement in this study.

### Declaration of generative AI and AI-assisted technologies in the writing process

### FIGURES AND LEGENDS

Figure 1

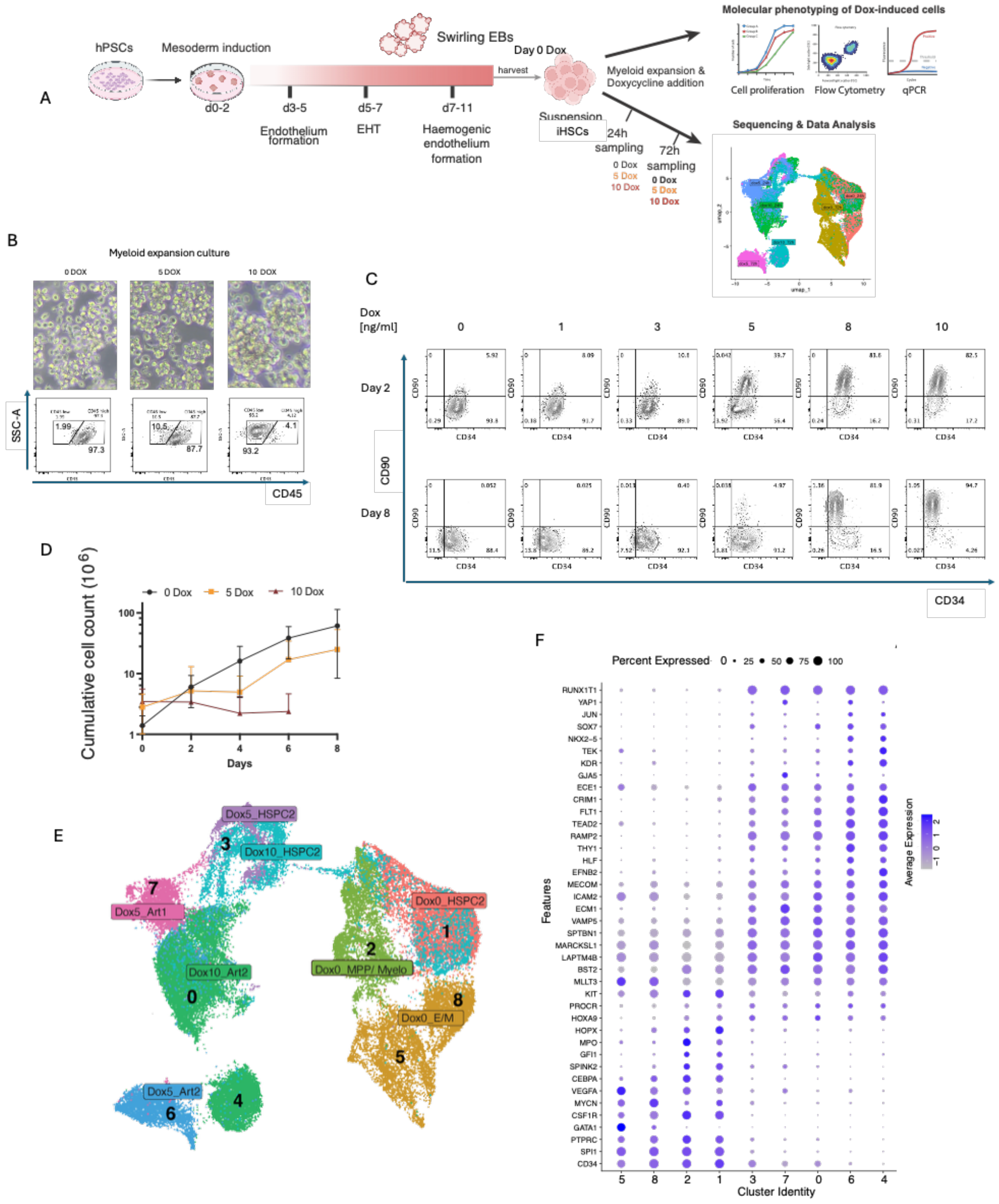

**Figure 1**

**Expression of RUNX1-ETO leads to dosage-dependent arterial patterning and a cell cycle block**

**A**: Schematic of the differentiation system showing the time course of hematopoietic differentiation from hPSCs to suspension iHSCs and further expansion after Doxycycline induction, followed by data analysis as depicted. Cells were expanded in rotating 6-well plates for up to 8 days and Doxycycline and growth medium were replenished every second day. Partially created using BioRender.com.

**B.** Brightfield images of RUNX1-ETO HPSC derived hematopoietic cells under no doxycycline (0 Dox), 5 ng/ml Dox and 10 ng/ml Dox conditions showing formation of cellular aggregates loss of CD45 expression (bottom panel).

**C.** Flow cytometric analysis of CD90 and CD34 surface marker expression at Day 2 and Day 8 of RUNX1-ETO induction using the indicated concentrations of Doxycycline (1,3,5,8,10 ng/ml) compared to uninduced cells (0 Dox).

**D.** Box and whisker plot showing cumulative cell counts following Dox-induction (0,5,10 ng/mL) conditions over 8 days. Error bars, mean ±s.e.m., n=3

**E.** UMAP representation of 8 Seurat clusters highlighting cell populations with annotated hematopoietic and non-hematopoietic identity as identified in Supplemental Figure 1 and by comparison with embryo data [26,27]. HSPC2: Hematopoietic stem and progenitor cells, embryonic cluster 2, Art1/2: Arterial cells, embryonic cluster 1 and 2, Myelo: Myeloid progenitor cells, E/M/M: Erythroid, megakaryocyte and mast cell progenitors.

**F.** Dot plot of expression levels of differentially expressed genes in Seurat clusters representing arterial, hemogenic endothelium and hematopoietic genes.

Fig 2

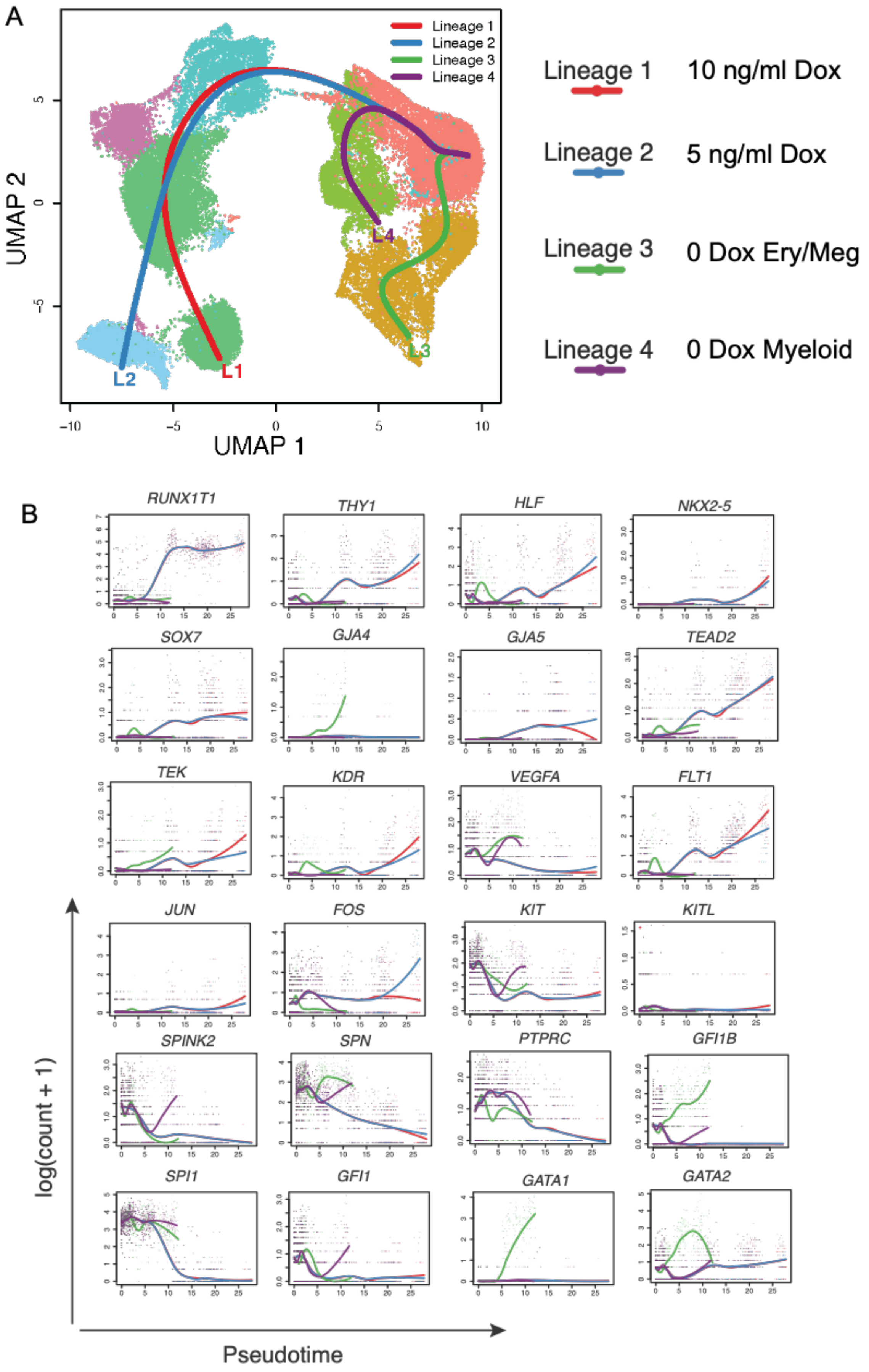
A
Lineage 1
Lineage 2
Lineage 3
Lineage 4
L1
L2
L3
L4
UMAP 1
UMAP 2
Lineage 1 10 ng/ml Dox
Lineage 2 5 ng/ml Dox
Lineage 3 0 Dox Ery/Meg
Lineage 4 0 Dox Myeloid
B
RUNX1T1
THY1
HLF
NKX2-5
SOX7
GJA4
GJA5
TEAD2
TEK
KDR
VEGFA
FLT1
JUN
FOS
KIT
KITL
SPINK2
SPN
PTPRC
GFI1B
SPI1
GFI1
GATA1
GATA2
log(count + 1)
Pseudotime

**Figure 2: Expression of RUNX1-ETO alters the developmental control of gene expression and the differentiation trajectory**

**A.:** Projection of the differentiation trajectory of cells in the four samples (0 Dox, 5 Dox and 10 Dox) on the 8 Seurat clusters highlighting cell populations with annotated hematopoietic and non-hematopoietic identity as in Fig 1E using trendSeq defining 4 lineages.

**B.:** (0 dox, 5 dox and 10 dox) on the Seurat clusters using tradeSeq defining 4 lineages. Lineage 1, 10 dox; Lineage 2, 5 dox; Lineages 3 and 4, 0 dox.

Figure 3

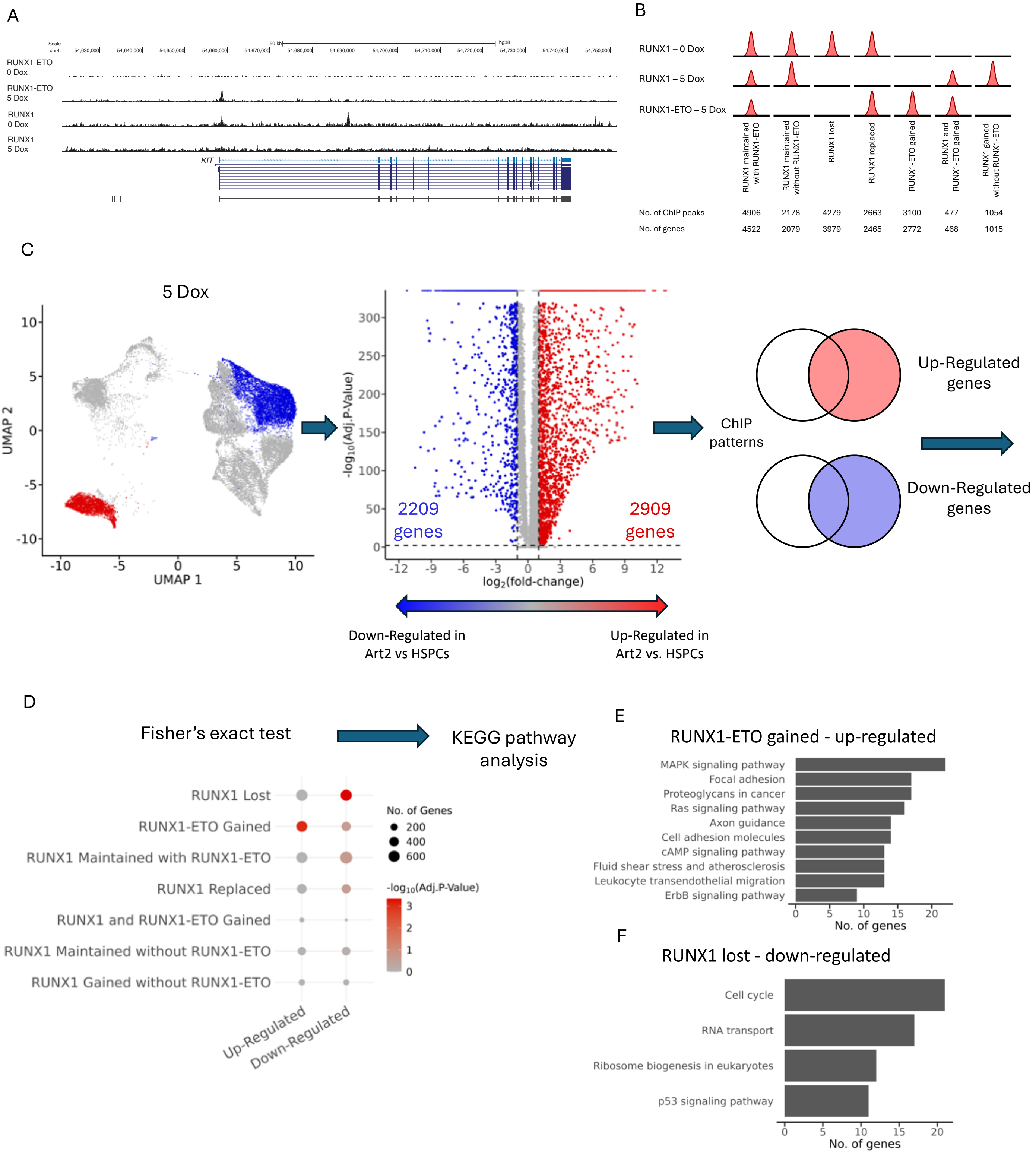
A
RUNX1-ETO 0 Dox
RUNX1-ETO 5 Dox
RUNX1 0 Dox
RUNX1 5 Dox
KIT
B
RUNX1 – 0 Dox
RUNX1 – 5 Dox
RUNX1-ETO – 5 Dox
RUNX1 maintained with RUNX1-ETO
RUNX1 maintained without RUNX1-ETO
RUNX1 lost
RUNX1 replaced
RUNX1-ETO gained
RUNX1 and RUNX1-ETO gained
RUNX1 gained without RUNX1-ETO
No. of ChIP peaks 4906 2178 4279 2663 3100 477 1054
No. of genes 4522 2079 3979 2465 2772 468 1015
C
5 Dox
UMAP 1
UMAP 2
-log10(Adj.P-Value)
log2(fold-change)
2209 genes
2909 genes
ChIP patterns
Up-Regulated genes
Down-Regulated genes
Down-Regulated in Art2 vs HSPCs
Up-Regulated in Art2 vs. HSPCs
D
Fisher's exact test
KEGG pathway analysis
RUNX1 Lost
RUNX1-ETO Gained
RUNX1 Maintained with RUNX1-ETO
RUNX1 Replaced
RUNX1 and RUNX1-ETO Gained
RUNX1 Maintained without RUNX1-ETO
RUNX1 Gained without RUNX1-ETO
Up-Regulated
Down-Regulated
No. of Genes
200
400
600
-log10(Adj.P-Value)
E
RUNX1-ETO gained - up-regulated
MAPK signaling pathway
Focal adhesion
Proteoglycans in cancer
Ras signaling pathway
Axon guidance
Cell adhesion molecules
cAMP signaling pathway
Fluid shear stress and atherosclerosis
Leukocyte transendothelial migration
ErbB signaling pathway
No. of genes
F
RUNX1 lost - down-regulated
Cell cycle
RNA transport
Ribosome biogenesis in eukaryotes
p53 signaling pathway
No. of genes

**Figure 3: RUNX1-ETO interferes with the RUNX1-driven gene regulatory network**

**A.** UCSC browser screenshots showing RUNX1 and RUNX1-ETO binding at the *KIT* locus in the presence or absence of 5 ng/ml Dox.

**B.** Depiction of seven different classes on RUNX1 and RUNX1-ETO binding patterns obtained from previously conducted ChiP experiments using the same cell line [18] from uninduced or 5 Dox induced HSPCs. Peaks were filtered against ATAC-Seq peaks. Shared peaks are depicted with half the height. The table below summarizes peak numbers per group. Genes associated with binding sites as determined by comparing accessible chromatin peaks to those known to be associated with their rightful promoter as determined by comparison with promoter-capture HiC data from human CD34+ cells [51] are listed in a bottom line.

**C.** Strategy of determining up- and down-regulated RUNX1 and RUNX1-ETO target genes in the 5 Dox-induced Art2 cluster (red) as compared to that of the uninduced HSPC2 cluster (blue). Left panel: UMAP of 0 and 5 dox cells highlighting the HSPC and Art2 cell populations at the start and end of the differentiation trajectory. Right: Volcano plot of genes which are differentially expressed between HSPC and Art2 cell populations in 0 and 5 dox cells. A positive fold-change indicates a gene was up-regulated in Art2 compared to HSPCs, and a negative value indicates a gene was down-regulated in Art2 compared to HSPCs. A gene was considered differentially expressed if it had a 2-fold change and an adjusted p-value less than 0.01. Right panel: Integration of differential gene expression values with the different patterns of RUNX1 and RUNX1-ETO binding as depicted in (B)

**D.** Results from a one-sided (right tailed) Fisher's exact test comparing the overlap of genes which were up and down-regulated in Art2 vs. HSPCs to ChIP binding groups depicted in Fig 2B. A significant p-value means that the number of genes which overlapped between groups was greater than could be expected by chance. Gene sets with a Benjamini-Hochberg corrected p-value $< 0.05$ was considered significant.

**E, F.** KEGG pathway enrichment of gene sets that were found to be significant, in this case up-regulated genes gaining RUNX1-ETO binding (E) and down-regulated genes that have lost RUNX1 binding. Lists of enriched genes can be found in (Supplemental dataset 5)

Figure 4

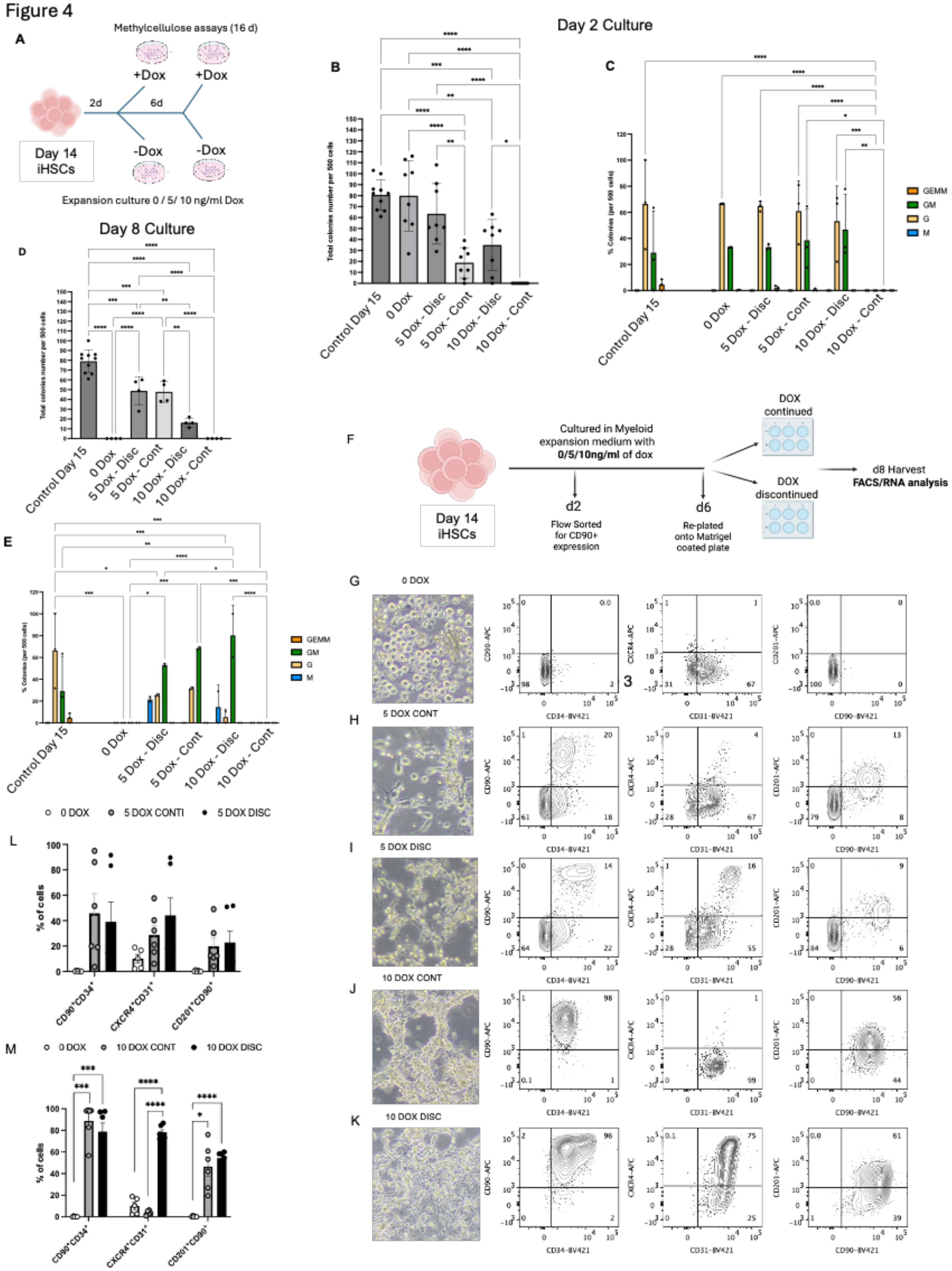

A
Methylcellulose assays (16 d)
+Dox
+Dox
2d
6d
Day 14
iHSCs
-Dox
-Dox
Expansion culture 0 / 5/ 10 ng/ml Dox
Day 2 Culture
B
C
Day 8 Culture
D
E
GEMM
GM
G
M
Control Day 15
0 Dox
5 Dox - Disc
5 Dox - Cont
10 Dox - Disc
10 Dox - Cont
F
Cultured in Myeloid expansion medium with 0/5/10ng/ml of dox
DOX continued
DOX discontinued
d8 Harvest
FACS/RNA analysis
d2
Flow Sorted for CD90+ expression
d6
Re-plated onto Matrigel coated plate
G
0 DOX
H
5 DOX CONT
I
5 DOX DISC
J
10 DOX CONT
K
10 DOX DISC
CD34-BV421
CD31-BV421
CD90-BV421
CD90-APC
CXCR4-APC
CD201-APC
L
0 DOX
5 DOX CONTI
5 DOX DISC
% of cells
M
0 DOX
10 DOX CONT
10 DOX DISC

**(L, M.)** Bar graphs of flow cytometry data for the indicated surface markers in cells treated as shown in (F). Error bars, mean ±s.e.m., n=6 experiments.

Figure 5

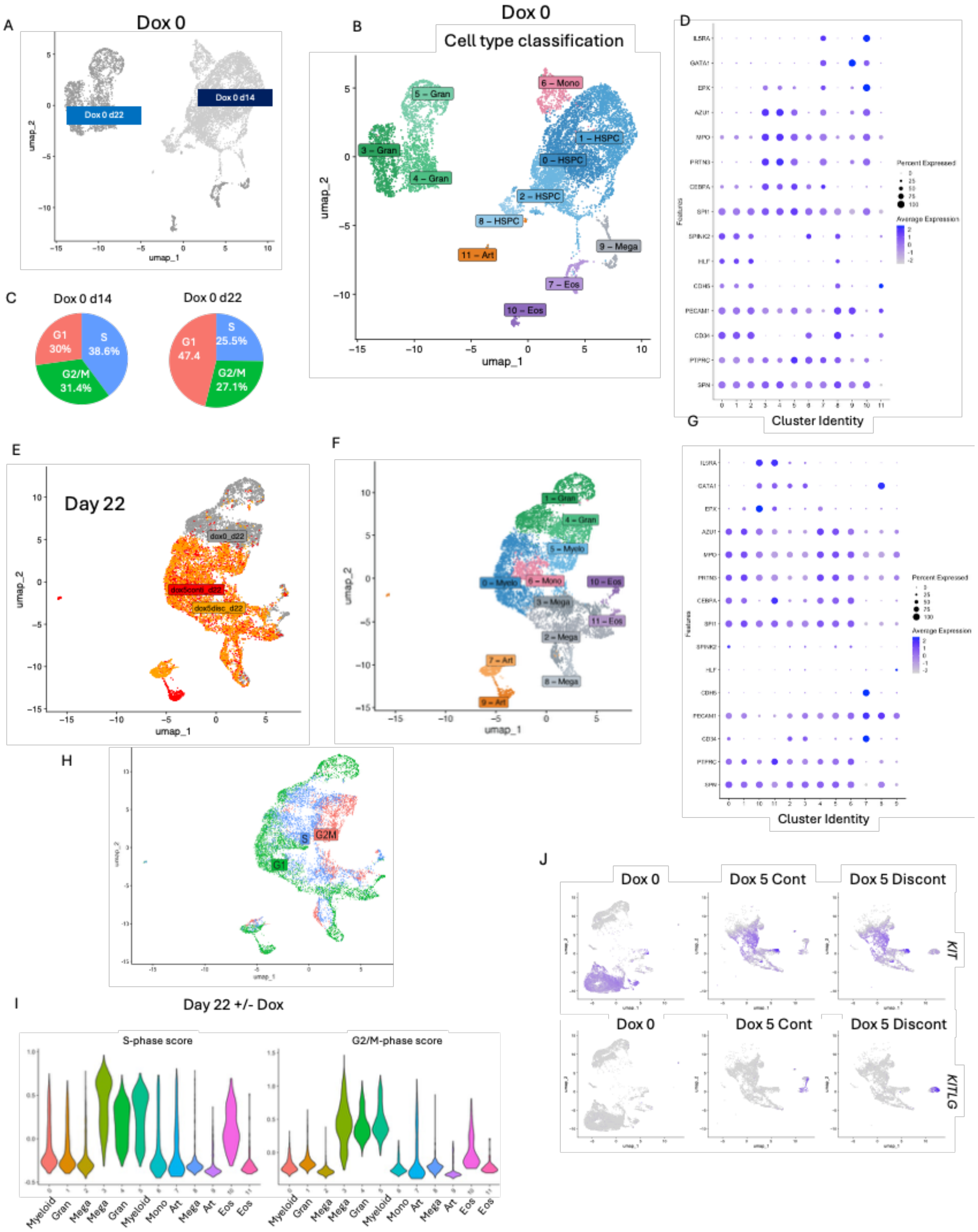
A
Dox 0
Dox 0 d22
Dox 0 d14
umap_1
umap_2
B
Dox 0
Cell type classification
5 – Gran
3 – Gran
4 – Gran
6 – Mono
1 – HSPC
0 – HSPC
2 – HSPC
8 – HSPC
11 – Art
9 – Mega
7 – Eos
10 – Eos
C
Dox 0 d14
G1 30%
S 38.6%
G2/M 31.4%
Dox 0 d22
G1 47.4
S 25.5%
G2/M 27.1%
D
IL5RA
GATA1
EPX
AZU1
MPO
PRTN3
CEBPA
SPI1
SPINK2
HLF
CDH5
PECAM1
CD34
PTPRC
SPN
Features
Percent Expressed
Average Expression
Cluster Identity
E
Day 22
dox0_d22
dox5cont_d22
dox5disc_d22
F
1 – Gran
4 – Gran
5 – Myelo
0 – Myelo
6 – Mono
10 – Eos
3 – Mega
11 – Eos
2 – Mega
7 – Art
8 – Mega
9 – Art
G
Cluster Identity
H
S
G2M
G1
I
Day 22 +/- Dox
S-phase score
G2/M-phase score
Myeloid
Gran
Mega
Mega
Gran
Myeloid
Mono
Art
Mega
Art
Eos
Eos
J
Dox 0
Dox 5 Cont
Dox 5 Discont
KIT
KITLG

**Figure 5: Endothelial reprogramming is irreversible, and endothelial culture supports the proliferation of RUNX1-ETO expressing hematopoietic cells**

**A.** UMAP representation of scRNA-Seq data from uninduced cells at Day 14 and Day 22.

**B.** Cell lineage annotation of uninduced cells at Day 14 and Day 22 to 11 clusters. HSPCs: Hematopoietic stem and progenitor cells; Art: Arterial endothelial cells; Gran: Granulocytic cells at different stages of maturation, Eos: Eosinophils, Mega: megakaryocytes, Mono: Monocytes.

**C.** Cell cycle status of uninduced Day 14 and Day 22 populations.

**D.** Dot plot of expression levels of differentially expressed genes in cells allocated to the Seurat clusters depicted in (B) representing stem cell and hematopoietic lineage genes.

**E.** UMAP representation of scRNA-Seq data from Day 22 cells from Matrigel cultures that were uninduced or induced with 5ng/ml Dox, followed by Dox continuation (dox5conti) or discontinuation (dox5disc) for the final 2 days of Matrigel culture (see Fig. 4F).

**F.** Cell lineage annotation of clusters of dox treated and untreated cells shown in (E) showing 9 clusters. Lineages as in (B).

**G.** Dot plot of expression levels of differentially expressed genes in cells allocated to Seurat clusters depicted in (F) representing hematopoietic genes.

**H.** UMAP representation of the cell cycle status of dox treated and untreated Day 22 cells from (E).

**I.** S-phase and G2M- scores of dox treated and untreated clusters of Day 22 cells from (F).

**J.** Feature plots from the same cell populations as in (F) separated by their induction status (Dox 0: No induction), continued induction with 5 ng/ml Dox (Dox 5 Cont) and after Dox withdrawal (Dox 5 Discont) showing the expression levels of *KIT* and *KITLG*

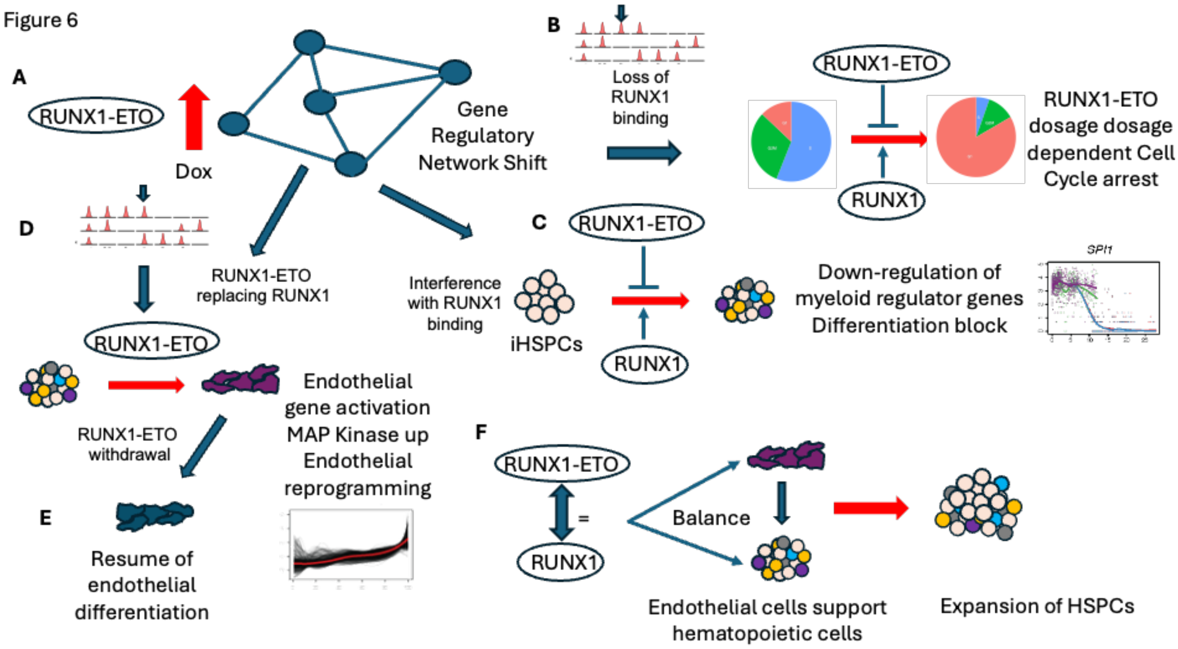


**Figure 6**

**Model of early events in RUNX1-ETO initiated leukemia development.**

**(A)** RUNX1-ETO expression the gene regulatory network of myeloid progenitor cells and interferes with RUNX1 binding**,**

**(B)** Deregulation of cell cycle genes is directly associated with RUNX1 loss and leads to a complete / partial cell cycle block**,**

**(C)** Down-regulation of myeloid regulator genes and differentiation block seen with all concentrations of RUNX1-ETO

**(D,E)** Rapid activation of endothelial gene expression program and MAP-Kinase pathway genes such as JUN directly associated with gain of RUNX1-ETO binding**, and (E)** Irreversible reprogramming of myeloid progenitor cells into endothelial cells after Dox withdrawal.

**(F)** Partial maintenance of hematopoietic gene expression program when RUNX1 and RUNX1-ETO expression levels are equal, but impediment of myeloid differentiation. Co-culture of hematopoietic and endothelial cells bypasses the block in proliferation of blood progenitors in the absence of hematopoietic growth factors.
